# Spastin tethers lipid droplets to peroxisomes and directs fatty acid trafficking through ESCRT-III

**DOI:** 10.1101/544023

**Authors:** Chi-Lun Chang, Aubrey V. Weigel, Maria S. Ioannou, H. Amalia Pasolli, C. Shan Xu, David R. Peale, Gleb Shtengel, Melanie Freeman, Harald F. Hess, Craig Blackstone, Jennifer Lippincott-Schwartz

## Abstract

Lipid droplets (LDs) are neutral lipid storage organelles that transfer lipids to various organelles including peroxisomes. Here, we show that the hereditary spastic paraplegia protein M1 Spastin, a membrane-bound AAA ATPase found on LDs, coordinates fatty acid (FA) trafficking from LDs to peroxisomes through two inter-related mechanisms. First, M1 Spastin forms a tethering complex with peroxisomal ABCD1 to promote LD-peroxisome contact formation. Second, M1 Spastin recruits the membrane-shaping ESCRT-III proteins IST1 and CHMP1B to LDs via its MIT domain to facilitate LD-to-peroxisome FA trafficking, possibly through IST1 and CHMP1B modifying LD membrane morphology. Furthermore, M1 Spastin, IST1 and CHMP1B are all required to relieve LDs of lipid peroxidation. M1 Spastin’s dual roles in tethering LDs to peroxisomes and in recruiting ESCRT-components to LD-peroxisome contact sites for FA trafficking may help explain the pathogenesis of diseases associated with defective FA metabolism in LDs and peroxisomes.

## Introduction

Subcellular organelles are highly collaborative in distributing metabolites, lipids, ions and proteins among themselves via vesicular and non-vesicular trafficking pathways. Non-vesicular pathways exploit the dense cytoplasmic packing of organelles to enable functional alliances through inter-organelle associations. Such associations are mediated by tethering proteins, which create so-called ‘contact sites’ for direct channeling of metabolites, lipids and ions between organelles (Wong et al., 2019). A major challenge has been to identify the diverse array of proteins involved in generating organelle contact sites and to understand how they mediate non-vesicular transport.

Among the organelles that engage in non-vesicular transport are lipid droplets (LDs) (Henne et al., 2018; Schuldiner and Bohnert, 2017). LDs function at the center of cell metabolism in many ways. They stockpile fatty acids (FAs) as neutral lipids, mainly triglycerides (TAG) and sterol esters, and release FAs as building materials for lipid synthesis and protein modification (Hashemi and Goodman, 2015; Pol et al., 2014; Walther et al., 2017). When nutrient availability is low, LDs transfer FAs to mitochondria and peroxisomes for β-oxidation, a crucial process that generates precursors for mitochondrial oxidative phosphorylation supporting energy production (Finn and Dice, 2006; Poirier et al., 2006). In addition, LDs transfer FAs to peroxisomes for various other steps in FA metabolism, including β-oxidation of very long chain FAs (VLCFAs), α-oxidation of branched chain FAs, bile acid and ether phospholipid synthesis, and docosahexaenoic acid (DHA) generation (Islinger et al., 2018; Lodhi and Semenkovich, 2014; Wanders, 2013). Aberrant FA metabolism in LDs is associated with severe physiological consequences, including neurological diseases and lipodystrophy (Kory et al., 2016; Welte, 2015). Defects in peroxisomal FA metabolism and biogenesis also lead to accumulation of LDs as demonstrated by patients with adrenoleukodystrophy (ALD) and Zellweger syndrome (Baes et al., 1997; Engelen et al., 2012; Schaumburg et al., 1972), suggesting a functional alliance between LDs and peroxisomes.

LDs reside in close proximity to both mitochondria and peroxisomes (Henne et al., 2018; Schuldiner and Bohnert, 2017). Contact sites between LDs and mitochondria and between LDs and peroxisomes are thought to facilitate direct channeling of FAs across these organelles’ boundaries and to prevent toxicity from free cytosolic FAs (Nguyen et al., 2017; Unger et al., 2010). While much attention has focused on characterizing contact sites between LDs and mitochondria, the machinery that creates LD-peroxisome contact sites and their functional significance in FA metabolism has remained a mystery.

Recently, possible protein candidates involved in LD-peroxisome contact sites and/or associated FA metabolism events have emerged from studies of the pathogenesis of hereditary spastic paraplegias (HSPs), a group of inherited neurological disorders with a prominent clinical feature of lower-extremity spasticity (Blackstone, 2018; Welte, 2015). Autosomal dominant mutations in the gene encoding Spastin are one of the most common causes of HSP and impact both LD and peroxisomal function. Depletion of Spastin in *Drosophila* and *C. elegans* leads to aberrant FA metabolism in LDs (Papadopoulos et al., 2015). In addition, HSP patient-derived olfactory neurosphere-derived cells with mutations in Spastin showed impaired peroxisome movement and distribution (Wali et al., 2016). This coincided with increased cellular lipid peroxidation and reduced energy production, possibly caused by defective FA trafficking to peroxisomes. Intriguingly, some ALD patients with mutations in a peroxisomal FA transporter ATP binding cassette subfamily D member 1 (*ABCD1*) also manifest neurologic symptoms, such as spasticity (Balicza et al., 2016; Koutsis et al., 2015; Lodhi and Semenkovich, 2014; Maris et al., 1995; Shaw-Smith et al., 2004; Zhan et al., 2013). Together, these findings raise the possibility that M1 Spastin and/or ABCD1 could function at LD-peroxisome contact sites to facilitate FA trafficking and FA metabolism in LDs and peroxisomes.

M1 Spastin is an isoform generated from the first translation initiation codon of Spastin (Claudiani et al., 2005). It contains an integral membrane hairpin motif in the N-terminal region that localizes it to LDs and other membrane compartments (Connell et al., 2009; Papadopoulos et al., 2015; Park et al., 2010; Reid et al., 2005). The shorter M87 Spastin isoform lacking this region resides primarily in the cytoplasm. M1 Spastin’s hairpin motif is followed by a microtubule interacting and trafficking (MIT) domain that selectively interacts with two endosomal sorting complexes required for transport (ESCRT)-III proteins, increased sodium tolerance 1 (IST1) and charged multivesicular body protein 1B (CHMP1B) (Agromayor et al., 2009; Reid et al., 2005; Renvoisé et al., 2010). Unlike most ESCRT-III components, IST1 and CHMP1B form external coats on positively-curved membrane (McCullough et al., 2015). M1 Spastin also contains a C-terminal microtubule binding (MTB) domain and an ATPase-Associated with diverse cellular Activities (AAA) ATPase domain implicated in remodeling microtubule networks (Roll-Mecak and Vale, 2008; White et al., 2007). Thus, M1 Spastin is a multi-modular ATPase combining the functions of LD association and ESCRT-III interaction. Nonetheless, a unifying mechanism that explains how different domains of Spastin operate together, specifically in FA metabolism at LDs and peroxisomes, is lacking.

Here, we provide evidence that M1 Spastin and ABCD1 form a tethering complex that draws LDs and peroxisomes together to facilitate LD-to-peroxisome FA trafficking. M1 Spastin promotes LD-peroxisome contact formation by inserting its N-terminal hairpin motif directly into the LD lipid monolayer and then using a peroxisome-interacting region to form a tethering complex with ABCD1 in peroxisomes. Furthermore, M1 Spastin controls LD-to-peroxisome FA trafficking by recruiting ESCRT-III proteins IST1 and CHMP1B to LDs. This indicates a novel role for ESCRT-III proteins in FA trafficking, possibly through modifying LD membrane morphology. Finally, we show that Spastin-mediated trafficking of FAs from LDs to peroxisomes is necessary for relieving LDs of peroxidated lipids. Together, these findings reveal a novel FA trafficking pathway orchestrated by M1 Spastin in conjunction with ESCRT-III and ABCD1 at LD-peroxisome contact sites, with relevance for understanding diseases involving mutations affecting Spastin and ABCD1.

## Results

### M1 Spastin promotes LD-peroxisome contact formation

To test a role of M1 Spastin in regulating the association between LDs and peroxisomes, we appended the fluorescence protein (FP) mApple to the N-terminus of M1 Spastin after mutating its second start codon to eliminate expression of M87 Spastin, which mostly resides in the cytoplasm (Figure 1A). When overexpressed in HeLa cells, mApple-M1 Spastin co-localized with BODIPY-labeled LDs, both under steady-state conditions and during LD biogenesis induced by oleic acid treatment (Figure 1B) (Papadopoulos et al., 2015). Using mEmerald-SKL to label peroxisomes, we found that peroxisomes normally have intermittent contacts with LDs (Figure 1C, top panel). Upon overexpression of M1 Spastin in these cells, however, a significant increase in association between LDs and peroxisomes occurred (Figure 1C, bottom panel), with the LD-associated peroxisomes frequently displaying an elongated morphology (Figure 1C, arrow heads in inset). Quantification of the fraction of LD signal overlapping with peroxisome signal revealed a significant increase in LD-peroxisome co-localization in M1 Spastin overexpressing cells as compared to control cells (Figure 1D). Immunostaining of endogenous peroxisomal proteins, including luminal protein catalase and membrane-associated protein PMP70, showed peroxisomes in close association with LDs decorated with mApple-M1-Spastin (Figure 1E). The enhancement of LD-peroxisome association by M1 Spastin overexpression was also observed in U-2 OS osteosarcoma cells and MRC-5 fibroblasts (Figures S1A and S1B).

**Figure 1.**
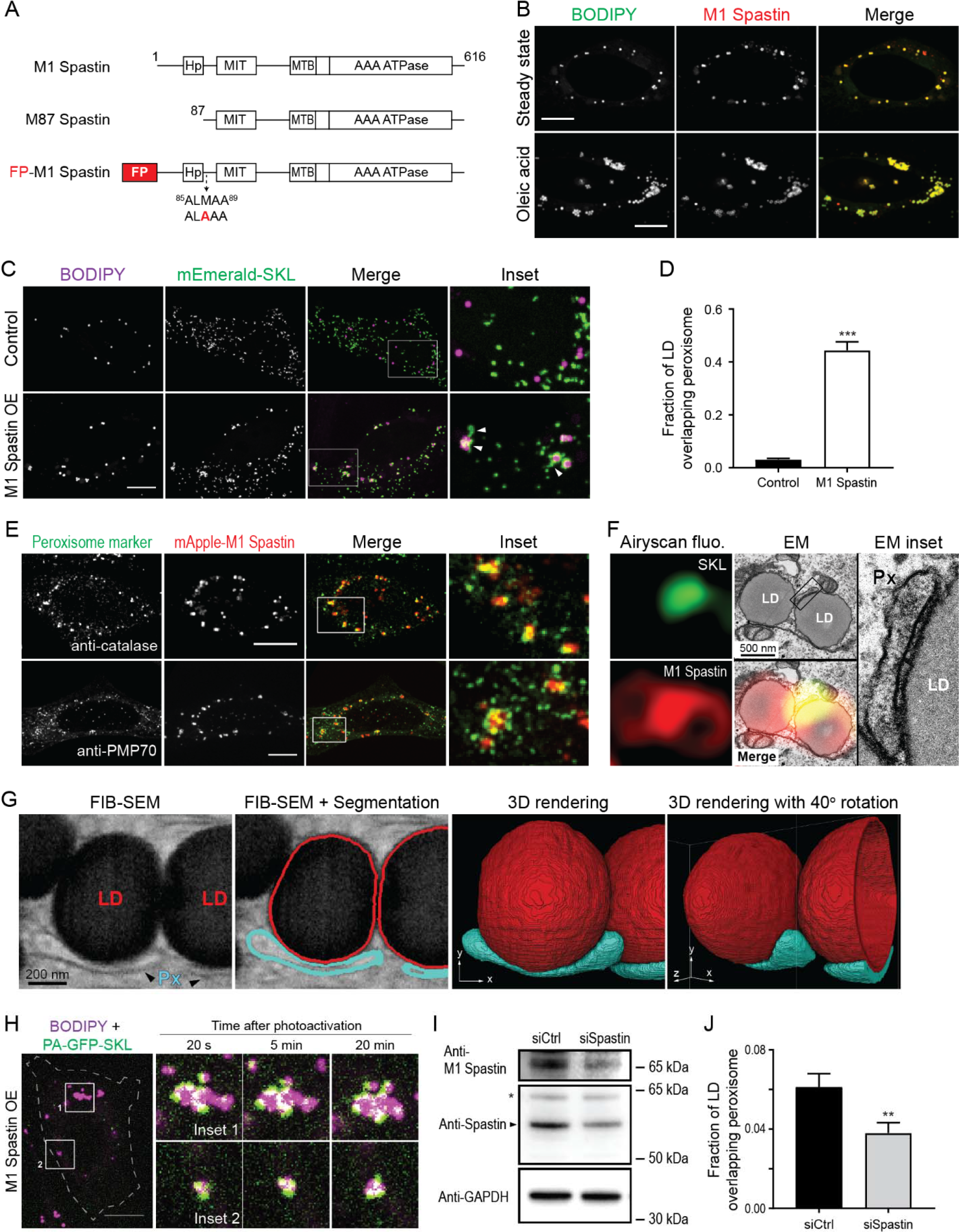
M1 Spastin promotes LD-peroxisome contact formation. (A) Diagrams of M1 Spastin, M87 Spastin, and fluorescence protein (FP)-tagged M1 Spastin construct. Amino acid number and protein domains are indicated. Hp, hairpin motif; MIT, microtubule interacting and trafficking domain; MTB, microtubule binding domain; AAA ATPase, ATPases Associated with diverse cellular Activities ATPase domain. The mutated residue of FP-M1 Spastin is labeled in red. (B) mApple-M1 Spastin co-localizes with BODIPY-493/503-labeled LDs in HeLa cells in steady state (top) or following 300 μM oleic acid treatment for 16 hours (bottom). Representative confocal images are shown. Scale bar, 10 μm. (C) Association between BODIPY-665/676-labeled LDs and mEmerald-SKL-labeled peroxisomes in control Hela cells or in Hela cells overexpressing (OE) mApple-M1 Spastin (not shown). Representative confocal maximal intensity projection (MIP) images are shown. White arrow heads indicate elongated peroxisomes associated with LDs. Scale bar, 10 μm. (D) Quantification of fraction of LD overlapping with peroxisomes as described in (C). Means ± SEM are shown (34-41 cells from 3 independent experiments). ***, p < 0.001. (E) Association between mApple-M1 Spastin and peroxisomes immunostained with catalase (top) and PMP70 (bottom) antibodies in Hela cells. Representative confocal MIP images are shown. Scale bar, 10 μm. (F) Correlative light electron microscopy (CLEM) images of a LD-peroxisomes contact site in mApple-M1 Spastin and mEmerald-SKL expressing HeLa cells following 15 μM oleic acid treatment for 16 hours. LD, lipid droplet; Px, peroxisome. (G) A single 8-nm FIB-SEM slice (left) at LD-peroxisome contacts in mApple-M1 expressing HeLa cells following 15 μM oleic acid treatment for 16 hours. LDs and peroxisomes (Px) were segmented (middle left) and reconstructed to reveal LD-peroxisome contacts in 3D (middle right and right). (H) Stable association between BODIPY-665/676-labeled LDs and peroxisomes labeled by PA-GFP-SKL in Hela cells overexpressing mApple-M1 Spastin monitored by confocal microscopy. Insets indicate photoactivation by 405 nm laser. Scale bar, 10 μm. (I) Spastin protein levels detected by Western blotting using anti-M1 Spastin (top) and anti-Spastin (middle) antibodies in HeLa cells transfected with siCtrl or siSpastin. GAPDH protein levels serve as a loading control (bottom). Arrow head and asterisk (middle panel) indicate M87 Spastin at expected molecular weight and a non-specific band, respectively. (J) Fraction of LD overlapping with peroxisomes in HeLa cells transfected with siCtrl or siSpastin. Means ± SEM are shown (25-28 cells from 3 independent experiments). **, p < 0.01.

We next used correlative light-electron microscopy (CLEM) to examine the nature of the association between LDs and peroxisomes in M1 Spastin expressing cells. In correlated Airyscan and transmission EM (TEM) images, elongated peroxisomes were visualized tightly attached to the surface of LDs (Figure 1F), indicating LD-peroxisome contacts. Increased electron density was seen at the contacts between the peroxisome and LD (Figure 1F, inset), prefiguring the existence of protein- and/or membrane-complexes that might serve to bridge the apposing organelles. We further applied focused ion beam-scanning EM (FIB-SEM) to examine LD-peroxisome contacts in 3D at 8-nm isotropic resolution in mApple-M1 Spastin expressing cells. Similar to conventional TEM, we observed elongated peroxisomes tightly associated with LDs in single FIB-SEM slices (Figure 1G). With the volumetric information provided by FIB-SEM, we could investigate the contact sites throughout the entirety of peroxisome and LD surfaces. The association between the peroxisome and LD spanned most of the length of the peroxisome. Interestingly, the contour of the entire peroxisomes mimicked that of the LDs. A tight association between the two organelles throughout the length of the peroxisome is seen, suggesting tethering sites with a large surface area (Figure 1G and movie S1).

Whether peroxisomes were stably tethered to or transiently interacting with LDs in M1 Spastin overexpressing cells was addressed by labeling peroxisomes with photoactivatable GFP attached to SKL (PA-GFP-SKL). Applying photoactivation to a few small regions containing LDs, a sub-population of peroxisomes was then tracked. The highlighted peroxisomes were found to be stably tethered to LDs during 20-min of imaging (Figure 1H), suggesting that M1 Spastin may function as a tether for LD-peroxisome contacts.

We next addressed whether endogenous M1 Spastin plays a role in LD-peroxisome contact formation. We first confirmed endogenous expression of M1 and M87 Spastin in HeLa cells transfected with control siRNA (siCtrl) by western blotting (Figure 1I). The expression of both Spastin isoforms was reduced in cells transfected with siRNA against Spastin (siSpastin). Despite the low occurrence of LD-peroxisome contacts in siCtrl-transfected cells, knockdown of Spastin led to a further reduction of LD-peroxisome co-localization (Figure 1J), indicating that endogenous M1 Spastin serves an important role in LD-peroxisome contact formation. Altogether, these results suggest that M1 Spastin promotes LD-peroxisome contact formation.

### M1 Spastin’s hairpin motif directly inserts into the lipid monolayer of LDs

To gain further insight into the role of M1 Spastin in LD-peroxisome tethering, we explored how M1 Spastin associates with LDs, focusing on its membrane-associated hydrophobic hairpin motif. Two synthetic constructs of M1 Spastin were generated that contain the first 92 amino acids of M1 Spastin appended with mApple FP, either at its N-terminus (FP-M1^1-92^) or its C-terminus (M1^1-92^-FP) (Figure 2A). Both mApple-M1^1-92^ and M1^1-92^-mApple labeled the endoplasmic reticulum (ER) and localized to LDs under both steady-state conditions and following oleic acid treatment (Figures S2A and S2B). We also observed co-localization of mApple-M1^1-92^ with GPAT4^152-208^, a membrane marker for nascent LDs (Wilfling et al., 2013) (Figure S2C). Therefore, M1^1-92^ appears to associate with LDs at different stages of LD biogenesis.

**Figure 2.**
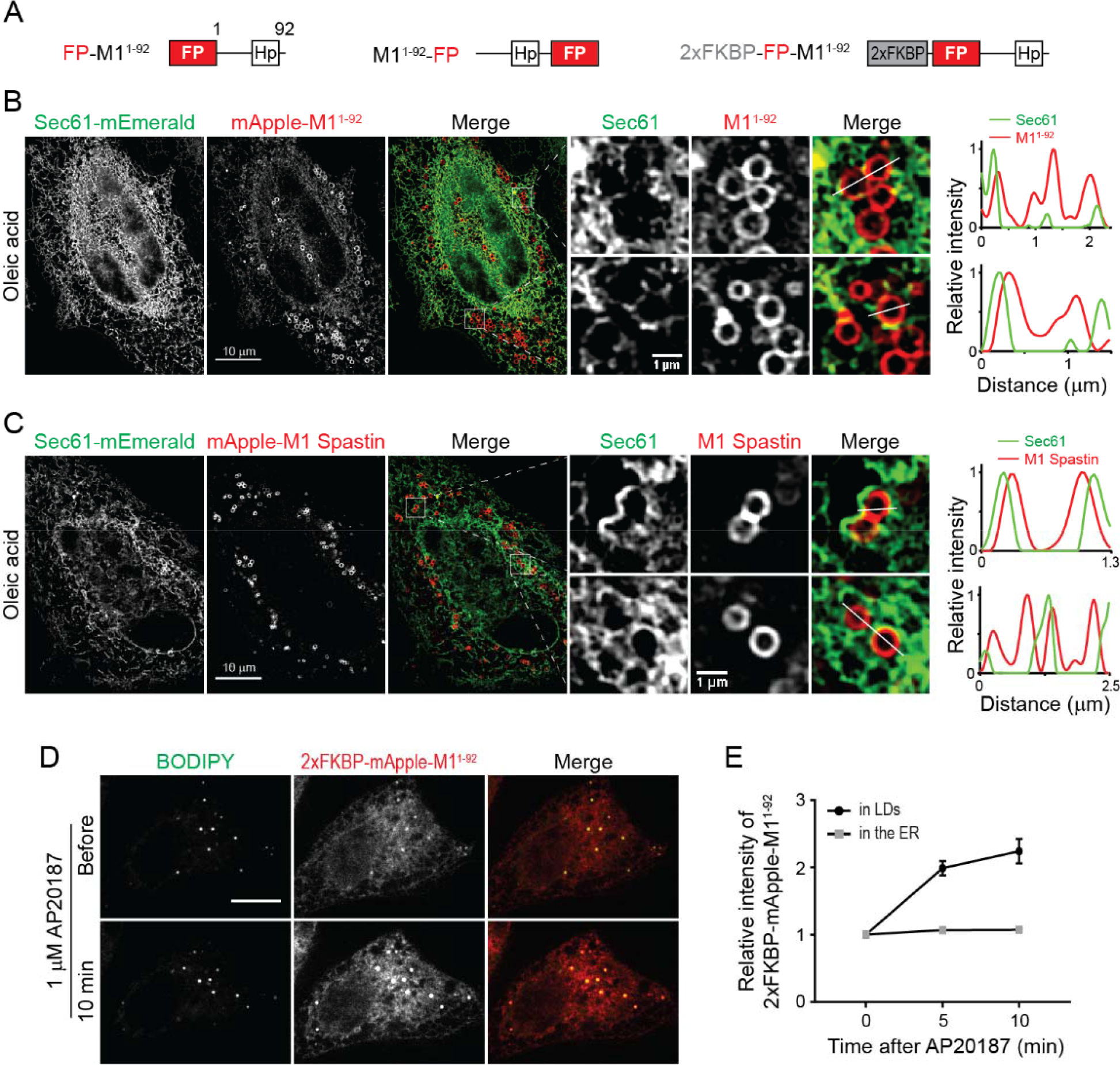
M1 Spastin’s hairpin motif directly inserts into the lipid monolayer of LDs. (A) Diagrams of synthetic constructs. Amino acid number and protein domains are indicated as in Figure 1A. 2xFKBP, tandem FKBP domain. (B) mApple-M1^1-92^ displays ER localization and ring structures enclosing LDs as monitored by structure illumination microscopy (SIM) in HeLa cells co-transfected with Sec61β-mEmerald following 300 μM oleic acid treatment for 16 hours. Line scans of relative intensity profile are shown (right panels). (C) mApple-M1 Spastin is localized to ring structures enclosing LDs as monitored by SIM in HeLa cells co-transfected with Sec61β-mEmerald following 300 μM oleic acid treatment for 16 hours. Line scans of relative intensity profile are shown (right panels). (D) Changes in intensity of 2xFKBP-mApple-M1^1-92^ in BODIPY-493/503-labeled LDs following 1 μM AP20187 treatment in Hela cells monitored by confocal microscopy. Scale bars, 10 μm. (E) Relative changes in intensity of 2xFKBP-mApple-M1^1-92^ in LDs and in the ER as described in (D). Means ± SEM are shown (12 cells from 2 independent experiments).

M1^1-92^ could co-localize with LDs by either directly inserting into the lipid monolayer of LDs or by an ER pool of M1^1-92^ wrapping tightly around LDs. To distinguish between these possibilities, we employed structured illumination microscopy (SIM), providing a two-fold increase in resolution compared to conventional confocal imaging, to visualize the ER labeled by Sec61β-mEmerald and mApple-M1^1-92^ in oleic acid-treated cells. SIM imaging of these cells coupled with line scan analysis revealed that ER tubules do not wrap around LDs labeled with mApple-M1^1-92^ (Figure 2B), supporting the idea that M1^1-92^ directly inserts into the lipid monolayer of LDs.

Like M1^1-92^, full-length M1 Spastin also localized to LDs by directly inserting into the LD monolayer, showing less co-localization with the ER marker Sec61β-mEmerald and greater enrichment on LDs than M1^1-92^ (Figure 2C). One reason for this difference could relate to full-length M1 Spastin’s ability to oligomerize into hexamers through the ATPase domain (Pantakani et al., 2008; Roll-Mecak and Vale, 2008), causing M1 Spastin to have higher affinity for LDs. To test this possibility, we inserted tandem FKBP (2xFKBP), a chemically-inducible oligomerization motif, to the N-terminus of mApple-M1^1-92^ to generate 2xFKBP-mApple-M1^1-92^ (Figure 2A), which should oligomerize in response to AP20187 treatment (Burnett et al., 2004). Before treatment, this protein localized to both LDs and the ER (Figure 2D). Shortly following AP20187 treatment, a 2-fold increase in the relative intensity of 2xFKBP-mApple-M1^1-92^ on LDs occurred (Figures 2D and 2E). These data suggest that oligomerization enhances the affinity of M1^1-92^ to LDs. Consistent with this, M1^1-92^ tagged with a dimeric red FP DsRed2 to enable protein oligomerization also showed primarily LD localization under steady-state conditions (Figure S2D). Together, these findings indicate that M1 Spastin directly inserts into the LD lipid monolayer via its N-terminal hairpin motif, with oligomerization enhancing this association.

### ATP hydrolysis is required for M1 Spastin to mediate LD-peroxisome contact formation

We next systematically surveyed the cytosolic regions of M1 Spastin to determine how it might induce LD-peroxisome contact formation. We found that M1 Spastin^Δ1-56^ lacking its N-terminal cytosolic region can still localize to LDs and drive LD-peroxisome contact formation (Figures S3A, S3B and S3E). Next, we generated a M1 Spastin^HFDD^ mutant, which has a defective MIT domain for ESCRT-III binding (Yang et al., 2008). Expression of the mutant protein revealed similar levels of LD localization and LD-peroxisome contacts as seen in cells expressing M1 Spastin (Figures S3C, S3D and S3E). Therefore, M1 Spastin’s N-terminal cytosolic region and MIT domain are not required for peroxisomal interaction.

To test whether M1 Spastin’s ATPase domain is important for mediating LD-peroxisome contacts, we generated ATPase deficient M1 Spastin mutants, including M1 Spastin^K388R^ and M1 Spastin^E442Q^ (Figure 3A). M1 Spastin^K388R^ and M1 Spastin^E442Q^ both localized to LDs (Figure S3F). M1 Spastin^E442Q^ additionally displayed a filamentous pattern. Notably, both mutations significantly abolished the ability of M1 Spastin to promote LD-peroxisome contact formation (Figures 3B and 3C), with greater reduction in the extent of LD-peroxisome contacts in M1 Spastin^E442Q^-expressing cells than in cells expressing M1 Spastin^K388R^ (Figure 3C). As the defect in ATPase activity is more pronounced in M1 Spastin^E442Q^ as compared to M1 Spastin^K388R^ (Evans et al., 2005), the results indicated that M1 Spastin’s ATPase activity is important for promoting LD-peroxisome contact formation.

**Figure 3.**
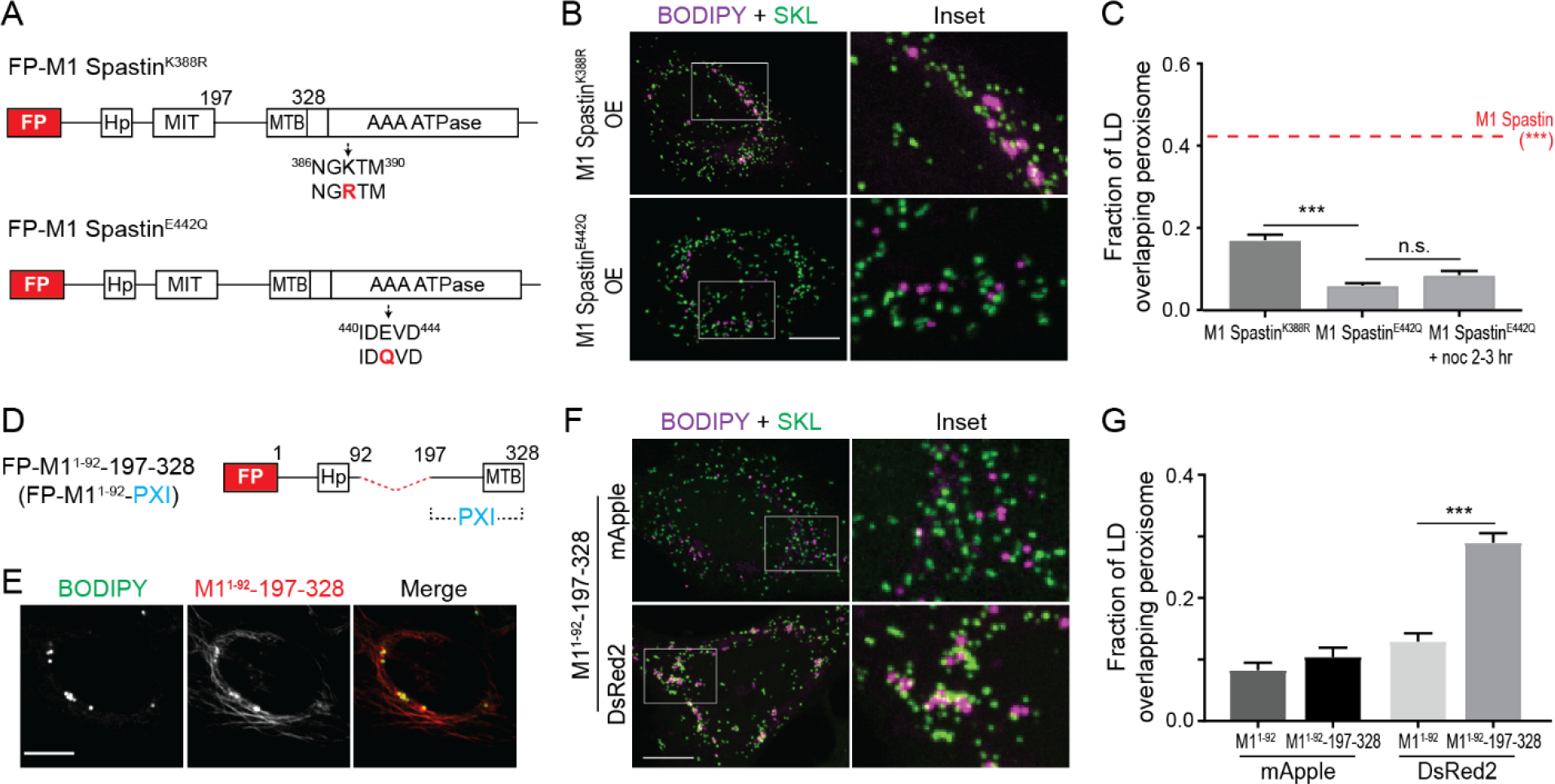
ATP hydrolysis is required for M1 Spastin to mediate LD-peroxisome contact formation via its PXI region. (A) Diagrams of FP-tagged M1 Spastin mutant constructs. Amino acid number and protein domains are indicated as Figure 1A. The mutated residues are labeled in red. (B) Localizations of BODIPY-665/676-labeled LDs and mEmerald-SKL-labeled peroxisomes in Hela cells overexpressing mApple-M1 Spastin^K388R^ (top) or mApple-M1 Spastin^E422Q^ (bottom). Representative confocal MIP images are shown. Scale bar, 10 μm. (C) Quantification of fraction of LD overlapping with peroxisomes as described in (B) and in cells overexpressing mApple-M1 Spastin (red dashed line). mApple-M1 Spastin^E422Q^-expressing cells were further placed on ice for 20 min and incubated with 10 μM nocodazole (noc) at 37°C for another 2-3 hours to disrupt microtubule network. Means ± SEM are shown (34-50 cells from at least 4 independent experiments). ***, p < 0.001. (D) Diagrams of a synthetic construct. The red dashed line indicates MIT domain deletion. PXI, peroxisome-interacting region. (E) Co-localization of BODIPY-665/676-labeled LDs and mApple-M1^1-92^-197-328 in HeLa cells. Representative confocal images are shown. Scale bar, 10 μm. (F) Association between BODIPY-665/676-labeled LDs labeled by and mEmerald-SKL-labeled peroxisomes in Hela cells overexpressing mApple-M1^1-92^-197-328 or DsRed2-M1^1-92^-197-328. Representative confocal MIP images are shown. Scale bar, 10 μm. (G) Quantification of fraction of LD overlapping with peroxisomes in HeLa cells overexpressing mApple-M1^1-92^, mApple-M1^1-92^-197-328, DsRed2-M1^1-92^ or DsRed2-M1^1-92^-197-328. Means ± SEM are shown (20-37 cells from at least 3 independent experiments). ***, p < 0.001.

The best-characterized function of M1 Spastin’s ATPase domain is microtubule severing (Roll-Mecak and Vale, 2008; White et al., 2007). To test whether microtubule disruption is important for LD-peroxisome contact formation, we depolymerized microtubules with nocodazole treatment to see if this could rescue the defect in LD-peroxisome contact formation in M1 Spastin^E442Q^-expressing cells. Upon depolymerization, no rescue was observed (Figure 3C), suggesting that microtubule severing is not critical for LD-peroxisome association induced by M1 Spastin. M1 Spastin’s ATPase domain, therefore, must be involved in some other aspect of LD-peroxisome interaction.

### Identification of a peroxisome-interacting (PXI) region on M1 Spastin

To investigate whether the cytosolic region between M1 Spastin’s MIT and ATPase domains plays a role in anchoring LDs to peroxisomes, we engineered a synthetic construct, M1^1-92^-197-328, that contains the amino acids 197-328 linked to M1^1-92^ for LD targeting (Figure 3D). This synthetic protein successfully targeted to BODIPY-labeled LDs (Figure 3E). The additional filamentous pattern observed could represent an ER pool of the protein bound to microtubules via its MTB domain (White et al., 2007).

Overexpression of mApple-M1^1-92^-197-328 did not promote LD-peroxisome contact formation (Figures 3F and 3G). However, M1^1-92^-197-328 expression led to significantly increased LD-peroxisome association when mApple was replaced with a dimeric red FP DsRed2 to enable protein oligomerization (Figures 3F and 3G). The results suggested, therefore, that amino acids 197-328 in M1 Spastin have affinity for peroxisomes, especially when the protein undergoes oligomerization. We have called this region of M1 Spastin PXI, for **p**ero**x**isome-**i**nteracting.

### ABCD1 forms a tethering complex with M1 Spastin via the PXI region

Some ALD patients with mutations in a peroxisome membrane protein ABCD1 manifest neurologic symptoms such as spasticity (Balicza et al., 2016; Koutsis et al., 2015; Maris et al., 1995; Shaw-Smith et al., 2004; Zhan et al., 2013), so we tested whether ABCD1 is required for LD-peroxisome contact formation by applying siRNA to knockdown ABCD1 (Figure S4A). The extent of LD-peroxisome contacts induced by M1 Spastin overexpression was significantly reduced in siABCD1-treated cells as compared to siCtrl-treated cells (Figures 4A and 4B). This suggested that ABCD1 is required for tethering LDs to peroxisomes.

**Figure 4.**
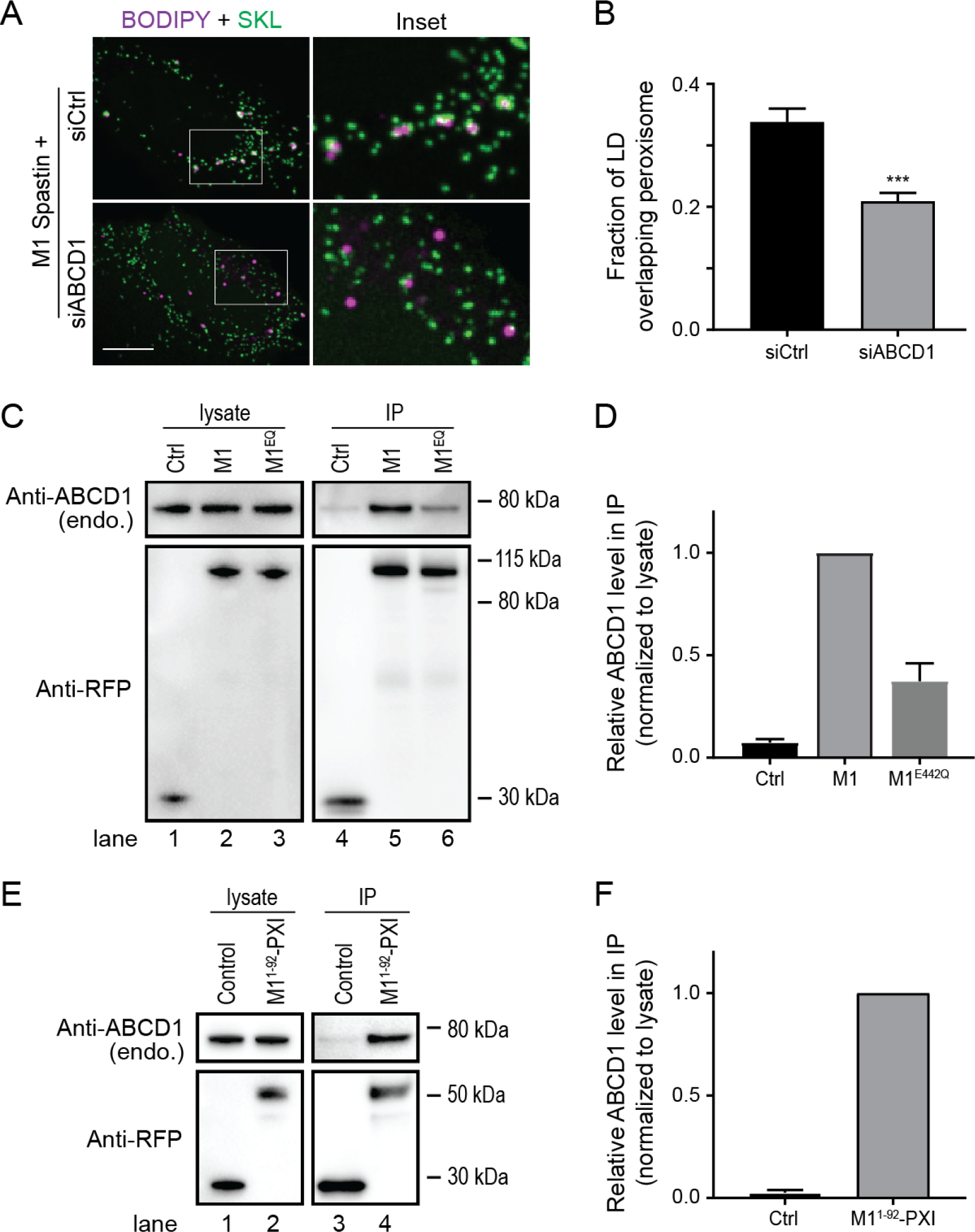
ABCD1 forms a tethering complex with M1 Spastin via the PXI region. (A) Localizations of BODIPY-665/676-labeled LDs and mEmerald-SKL-labeled peroxisomes in mApple-M1 Spastin-expressing Hela cells co-transfected with siCtrl or siABCD1. Representative confocal MIP images are shown. Scale bar, 10 μm. (B) Quantification of fraction of LD overlapping with peroxisomes as described in (A). Means ± SEM are shown (39-44 cells from 4 independent experiments). ***, p < 0.001. (C) Immunoprecipitation (IP) of ABCD1 in Hela cells transfected with mApple-C1 vector (Ctrl), mApple-M1 Spastin (M1), or mApple-M1 Spastin^E442Q^ (M1^EQ^). Protein levels of endogenous ABCD1 and overexpressed constructs in cell lysates and IP were assessed by Western blotting using antibodies against ABCD1 and RFP. (D) Quantification of relative ABCD1 level in the IP as described in (C). The value of M1 was set as 1. Means ± SEM from 3 independent IP experiments are shown. (E) IP of ABCD1 in Hela cells transfected with mApple-C1 vector (Control) and mApple-M1^1-92^-PXI. Protein levels of endogenous ABCD1 and overexpressed constructs in cell lysates and IP were assessed by Western blotting using antibodies against ABCD1 and RFP. (F) Quantification of relative ABCD1 level in the IP as described in (E). The value of mApple-M1^1-92^-PXI was set as 1. Means ± SEM from 2 independent IP experiments are shown.

To test if ABCD1 forms a complex with M1 Spastin for LD-peroxisome tethering, we performed immunoprecipitation (IP) experiments. Endogenous ABCD1 co-immunoprecipitated with mApple-M1 Spastin (M1) but not with mApple FP (Figure 4C, lane 4 and 5; Figure 4D). M1 Spastin seemed to associate selectively with ABCD1, since we did not detect comparable levels of ACBD5, another peroxisomal membrane protein, in our IP experiments (Figures S4B and S4C). The ability of M1 Spastin to associate with ABCD1 was reduced when an E442Q mutation was introduced into Spastin (M1^EQ^) (Figure 4C, lanes 5 and 6; Figure 4D), suggesting that ATPase activity is important for forming the M1 Spastin-ABCD1 tethering complex. We further found that endogenous ABCD1 co-immunoprecipitated with overexpressed M1^1-92^-197-328, namely M1^1-92^-PXI, since it contained the PXI region residues 197-328 (Figures 4E and 4F). This indicated that the PXI region of M1 Spastin is sufficient to mediate complex formation with ABCD1. Altogether, these data reveal that ABCD1 forms a tethering complex with the PXI region of M1 Spastin.

### M1 Spastin controls fatty acid trafficking from LDs to peroxisomes

Since ABCD1 is a FA transporter on peroxisomes (Lodhi and Semenkovich, 2014) and appears to form a tethering complex with M1 Spastin at LD-peroxisome contacts, we wondered whether FA trafficking from LDs to peroxisomes could be regulated by M1 Spastin. To monitor LD-to-peroxisome FA trafficking in cells, we used the fluorescence FA analog NBD-C12 in a FA pulse-chase assay in which cells were pulse-labeled by incubating with trace amounts of NBD-C12 for 24 hours and then chased in label-free medium (Figure S5A). Visualization of cells immediately after the pulse labeling revealed that NBD-C12 had incorporated into BODIPY-labeled LDs (Figure S5B, upper panels). Upon chasing for 18 hours, a significant pool of NBD-C12 had now redistributed into peroxisomes (Figure S5B, middle panels), indicating LD-to-peroxisome FA trafficking. Incubation with a pan-lipase inhibitor DEUP during the chase abolished the redistribution of NBD-C12 from LDs to peroxisomes (Figure S5B, bottom panels). The DEUP-treated cells showed substantial accumulation of NBD-C12 in LDs and a moderate reduction of NBD-C12 in peroxisomes as compared to non-treated cells (Figure S5C). This indicated that release of NBD-C12 from neutral lipids via lipase activities is required for NBD-C12 redistribution from LDs to peroxisomes. Together, these results indicated that NBD-C12 can be used to monitor lipase-sensitive FA trafficking from LDs to peroxisomes.

To test if endogenous Spastin is required for transfer of FAs from LDs to peroxisomes, we performed the NBD-C12 pulse-chase assay in Spastin knockdown cells (Figure 5A). NBD-C12 signal accumulated in LDs in siSpastin cells, but weaker NBD-C12 signal in peroxisomes was observed relative to siCtrl-treated cells (Figures 5B and 5C). This suggested that endogenous Spastin is important for FA trafficking from LDs to peroxisomes.

**Figure 5.**
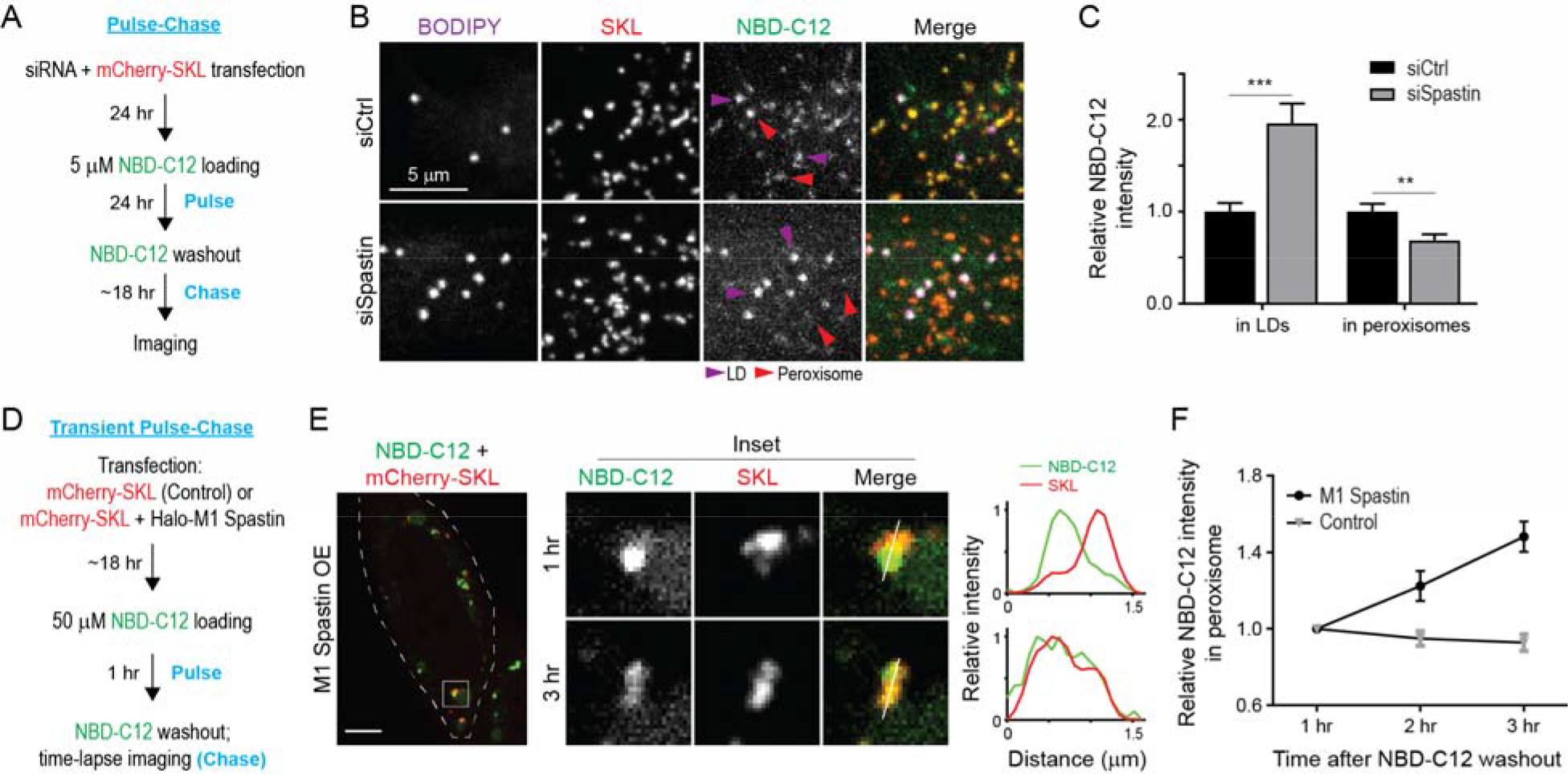
M1 Spastin controls LD-to-peroxisome FA trafficking. (A) Schematic diagram representing NBD-C12 FA pulse-chase assay in cells transfected with siRNAs and mCherry-SKL. (B) Distribution of NBD-C12 in BODIPY-665/676-labeled LDs and mCherry-SKL-labeled peroxisomes in HeLa cells transfected with siCtrl or siSpastin. Magenta and red arrow heads indicate LDs and peroxisomes, respectively. Representative confocal MIP images are shown. (C) Relative NBC-C12 intensity in LDs or peroxisomes as described in (B). Means ± SEM are shown (26-30 cells from 3 independent experiments). **, p < 0.01; ***, p < 0.001. (D) Schematic diagram representing transient NBD-C12 FA pulse-chase assay in cells transfected with mCherry-SKL alone or mCherry-SKL and Halo-M1 Spastin. (E) Redistribution of NBD-C12 to peroxisomes in Halo-M1 Spastin overexpressing HeLa cells monitored by confocal microscopy. Line scans of relative intensity profile are shown (right panels). Representative confocal mages are shown. Scale bar, 10 μm. (F) Relative NBC-C12 intensity in peroxisomes as described in (D and E). Means ± SEM are shown (12-13 cells from 3 independent experiments).

We next examined whether M1 Spastin overexpression could enhance LD-to-peroxisome FA trafficking. For this analysis, we modified the above pulse-chase assay by pulsing cells with NBD-C12 for only 1 hour before chasing to monitor dynamic FA distribution to peroxisomes (Figure 5D). NBD-C12 was successfully incorporated into LDs after pulse labeling as its signal accumulated in round-shaped structures (Figure 5E, left panel) and co-localized with M1 Spastin (not shown). After 1 hour chase, most NBD-C12 fluorescence remained in LDs with minimal levels of NBD-C12 transferred to nearby peroxisomes, as shown by line scan analysis of intensity profiles from NBD-C12 and SKL signals (Figure 5E, middle and right panels). After 3 hours of chase, however, significant levels of NBD-C12 had accumulated in peroxisomes, with most NBD-C12 labeling in LDs gone. Quantification showed a steady increase in NBD-C12 intensity in peroxisomes in M1 Spastin-overexpressing cells, while no increase was observed in control cells (Figure 5F). These data indicated that M1 Spastin overexpression enhances FA trafficking from LDs to peroxisomes.

### M1 Spastin recruits ESCRT-III proteins IST1 and CHMP1B to LDs

M1 Spastin has an MIT domain that interacts with the membrane shaping ESCRT-III proteins IST1 and CHMP1B (Agromayor et al., 2009; McCullough et al., 2015; Reid et al., 2005; Renvoisé et al., 2010). We tested, therefore, whether M1 Spastin could recruit IST1 and CHMP1B onto LDs. Consistent with this notion, FP-tagged IST1 and CHMP1B labeled bright loci that co-localized with mApple-M1 Spastin (Figure 6A). The IST1- and CHMP1B-labeled loci disappeared in cells co-expressing the M1 Spastin^HFDD^ mutant with a mutant MIT domain unable to bind ESCRT-III (Figures 6B). Examination of IST1-labeled loci by Airyscan microscopy and line-scan analysis revealed that IST1 was restricted to the surface of round-shaped LDs and not to nearby SKL-labeled peroxisomes (Figure 6C). These results indicated that IST1 and CHMP1B are recruited to LDs by M1 Spastin’s MIT domain.

**Figure 6.**
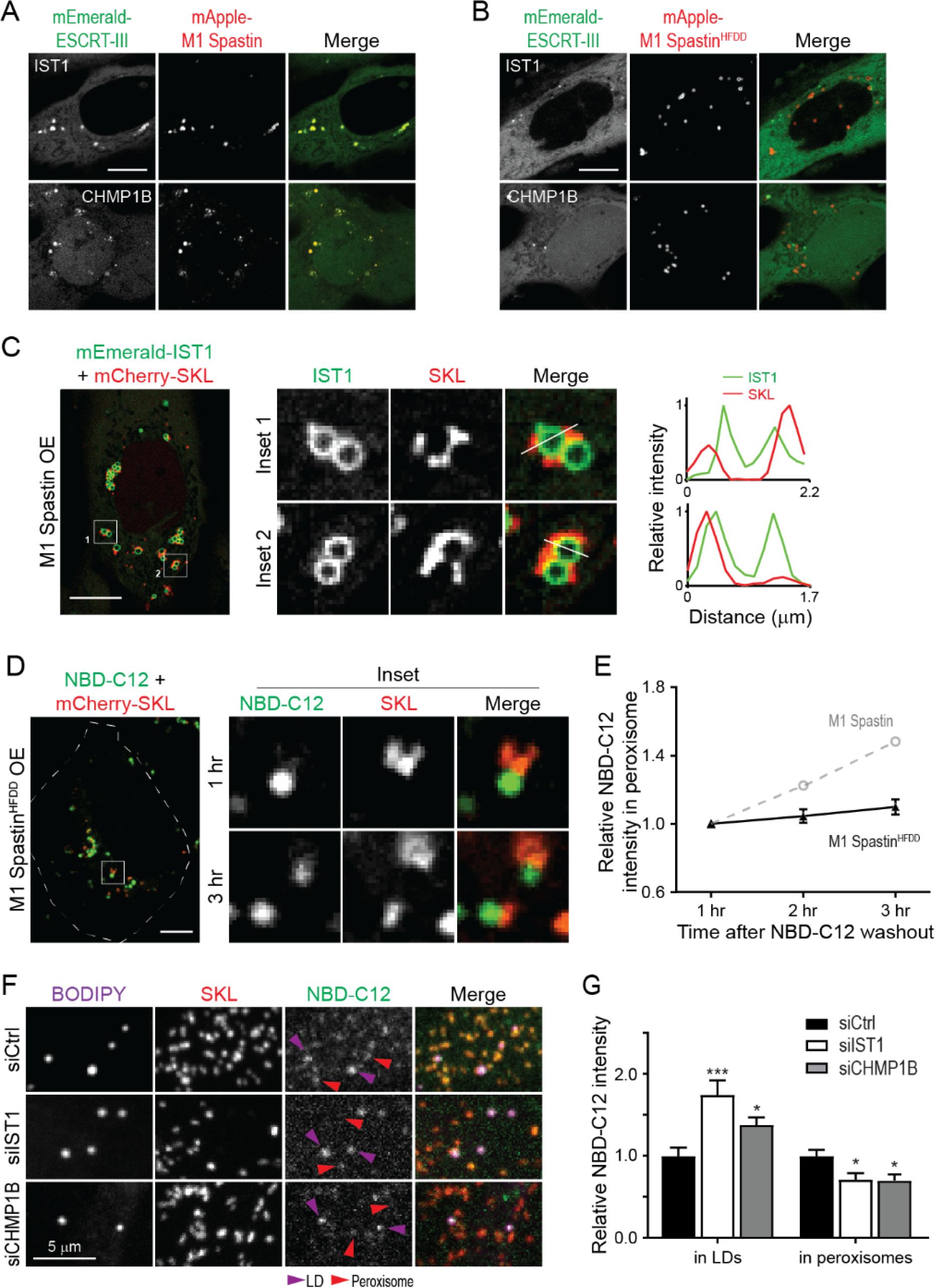
M1 Spastin recruits IST1 and CHMP1B to LDs to support LD-to-peroxisome FA trafficking. (A and B) Subcellular localization of mEmerald-IST1 (top panels) and mEmerald-CHMP1B (bottom panels) in Hela cells overexpressing mApple-M1 Spastin (A) or mApple-M1 Spastin^HFDD^ (B) monitored by confocal microscopy. Representative confocal images are shown. Scale bars, 10 μm. (C) mEmerald-IST1 is recruited to LDs but not to mCherry-SKL-labeled peroxisomes in HeLa cells co-expressing Halo-M1 Spastin monitored by Airyscan microscopy. Line scans of relative intensity profile are shown (right panels). Scale bar, 10 μm. (D) Distribution of NBD-C12 in Halo-M1 Spastin^HFDD^ and mCherry-SKL overexpressing HeLa cells monitored by confocal microscopy. Scale bar, 10 μm. (E) Relative NBC-C12 intensity in peroxisomes as described in (D). Means ± SEM are shown (19 cells from 3 independent experiments). Gray dashed line is a replica from Figure 5F. (F) Distribution of NBD-C12 in BODIPY-665/676-labeled LDs and mCherry-labeled peroxisomes in HeLa cells transfected with siCtrl, siIST1, or siCHMP1B. Representative confocal MIP images are shown. (G) Relative NBC-C12 intensity in LDs or peroxisomes as described in (F). Means ± SEM are shown (25-31 cells from 3 independent experiments). *, p < 0.05; ***, p < 0.001 as compared to siCtrl.

### IST1 and CHMP1B are required for M1 Spastin-regulated FA trafficking

Membrane curvature is known to contribute to lipid transfer between membranes (Lev, 2012). Since the ESCRT-III proteins IST1 and CHMP1B are curvature generators, we examined if recruitment of IST1 and CHMP1B to LDs by M1 Spastin is important for FA trafficking. To address this, we used the 1-hour FA pulse-chase labeling assay described in Figure 5D to assess LD-to-peroxisome FA trafficking in cells expressing the M1 Spastin^HFDD^ mutant unable to recruit ESCRT-III proteins.

Similar to M1 Spastin overexpressing cells, NBD-C12 successfully incorporated into LDs after 1 hour of labeling in cells overexpressing M1 Spastin^HFDD^ (Figure 6D). However, only a small amount of NBD-C12 appeared in peroxisomes after a 3-hour chase in these cells (Figures 6D and 6E), even though M1 Spastin^HFDD^ could enhance LD-peroxisome contact formation to the same extent as M1 Spastin (Figures S3C and S3D). As M1 Spastin^HFDD^ lacks a functional MIT domain to recruit ESCRT-III proteins to the surface of LDs, the data support the idea that ESCRT-III proteins are involved in FA trafficking from LDs to peroxisomes.

Consistently, knockdown of IST1 or CHMP1B also led to a reduction of peroxisomal NBD-C12 labeling as compared with siCtrl cells in experiments employing the NBD-C12 pulse-chase assay described in Figure 5A (Figures 6F, 6G, S6A, and S6B). Reduction of peroxisomal NBD-C12 labeling coincided with NBD-C12 accumulation in LDs in siIST1 or siCHMP1B cells, suggesting impaired FA release from LDs. Overall, these results demonstrated that M1 Spastin controls LD-to-peroxisome FA trafficking by recruiting ESCRT-III proteins IST1 and CHMP1B to LDs.

### The pathogenic Spastin^K388R^ mutant disrupts LD-peroxisome contact formation leading to peroxidated lipid accumulation in LDs

Peroxisomes are essential for relieving oxidative stress and maintaining redox homeostasis (Wanders, 2013). We wondered if FA trafficking at LD-peroxisome contacts may influence the extent of lipid peroxidation in LDs. We designed an imaging-based approach using LD-localized BODIPY-C11 (Figure S7A), a FA analog that when peroxidated shifts its emission from 590 nm to 510 nm, as a sensor for measuring peroxidated lipids in LDs (Figure 7A). We applied cumene hydroperoxide (cumyl-OOH) treatment to induce lipid peroxidation (van der Kraaij et al., 1990). In control cells, we observed an increase in lipid peroxidation within LDs upon cumyl-OOH treatment (Figures 7A and 7B). Overexpression of M1 Spastin significantly reduced the levels of LD peroxidation caused by cumyl-OOH treatment. Given our data showing M1 Spastin’s role in mediating LD-to-peroxisome FA trafficking, these findings suggested the reduced LD peroxidation levels in M1 Spastin expressing cells was due to peroxidated FAs being transferred to peroxisomes.

**Figure 7.**
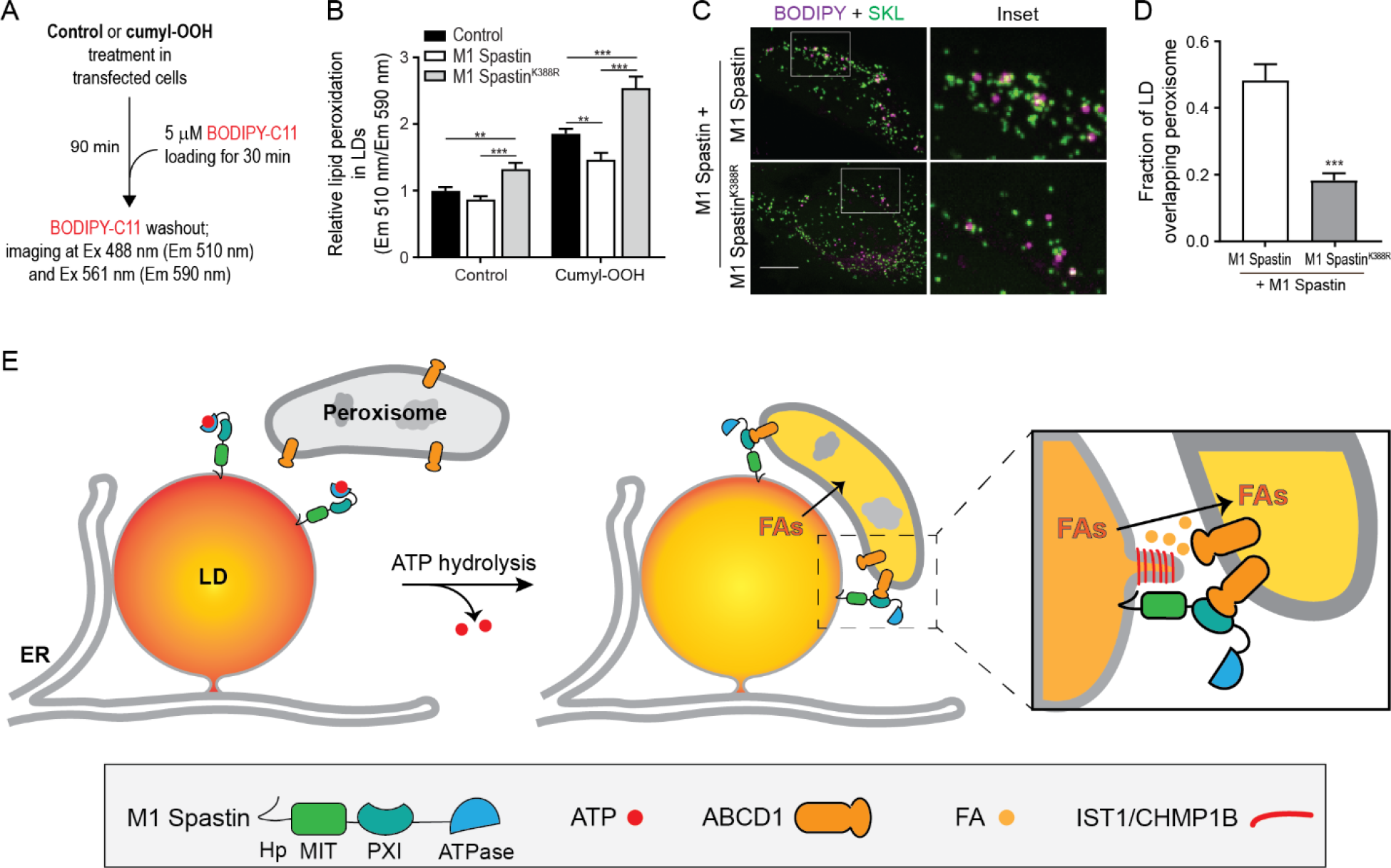
The pathogenic Spastin^K388R^ mutant disrupts LD-peroxisome contact formation leading to peroxidated lipid accumulation in LDs. (A) Schematic diagram representing lipid peroxidation assay using BODIPY-C11 to monitor LD peroxidation in the absence orpresence of cumene hydroperoxide (cumyl-OOH). (B) Relative lipid peroxidation in LDs as indicated by emission ratio at 510 nm over 590 nm in control HeLa cells or in those expressing Halo-M1 Spastin or Halo-M1 Spastin^K388R^ in the absence or presence of 50 μM cumyl-OOH. Means ± SEM are shown (32-56 cells from at least 3 independent experiments). **, p < 0.01; ***, p < 0.001. (C) Association between BODIPY-665/676-labeled LDs and mEmerald-SKL-labeled peroxisomes in Hela cells co-expressing mApple-M1 Spastin and mTagBFP2-M1 Spastin or mTagBFP2-M1 Spastin^K388R^. Representative confocal MIP images are shown. Scale bar, 10 μm. (D) Quantification of fraction of LD overlapping with peroxisomes as described in (C). Means ± SEM are shown (13-17 cells from 2 independent experiments). ***, p < 0.001. (E) Model of M1 Spastin in conjunction with ABCD1, IST1 and CHMP1B regulating LD-peroxisome contact formation and LD-to-peroxisome FA trafficking.

We next examined how the HSP pathogenic mutant M1 Spastin^K388R^ impacts LD-to-peroxisome FA trafficking and lipid peroxidation in LDs. M1 Spastin^K388R^ was incapable of promoting LD-peroxisome contact formation (Figures 3B and 3C) and no LD-to-peroxisome FA trafficking was observed in the transient NBD-C12 pulse-chase assay in cells overexpressing M1 Spastin^K388R^ (Figures S7B and S7C). M1-Spastin^K388R^ expression also led to a significant increase in LD peroxidation even in resting cells. This suggested that M1 Spastin^K388R^ disrupts the endogenous LD-to-peroxisome FA trafficking pathway. Consistent with this, following cumyl-OOH treatment, M1 Spastin^K388R^-expressing cells showed a nearly 2-fold increase in LD peroxidation as compared to M1 Spastin-expressing cells.

The K388R mutation in Spastin is an autosomal dominant mutation for HSP (Blackstone, 2018), suggesting that Spastin^K388R^ might have a dominant negative effect on M1 Spastin-mediated LD-peroxisome contact formation. Consistent with this possibility, we found a significant reduction in LD-peroxisome contacts in M1 Spastin expressing cells co-transfected with M1 Spastin^K388R^ or M87 Spastin^K388R^ (Figures 7C, 7D, S7D and S7E). This dominant negative effect on LD-peroxisome tethering might explain the elevated lipid peroxidation in LDs that we observed in M1 Spastin^K388R^ overexpressing cells.

## Discussion

In this paper, we identify a protein complex that tethers LDs to peroxisomes and show that the complex is critical for FA trafficking between these organelles. Comprised of LD-associated M1 Spastin and peroxisomal FA transporter ABCD1, the complex works in conjunction with ESCRT-III proteins IST1 and CHMP1B in delivering FAs from LDs to peroxisomes. Below, we discuss the basis for these findings, their significance for understanding the mechanics of inter-organelle FA trafficking, and their relevance to the molecular basis of certain types of HSP and diseases involved in defective peroxisome functions.

Frequent LD-peroxisome contact sites have been shown in professional FA-handling cells, such as adipocytes and hepatocytes, as well as other cell types including yeast and plant cells (Binns et al., 2006; Hayashi et al., 2001; Novikoff and Novikoff, 1973; Novikoff et al., 1980; Novikoff et al., 1973; Schrader, 2001; Thazar-Poulot et al., 2015). These studies demonstrated that LD-peroxisome contacts are evolutionarily-conserved inter-organelle structures and prompted speculation that directed transfer of FAs occurs at these contact sites. It has been a mystery, however, how peroxisomes and LDs coordinate with each other to facilitate this metabolic crosstalk.

We reasoned that some type of protein complex should tether LDs and peroxisomes together and modulate their FA trafficking. Seeking such a LD-peroxisome tethering complex, we found that M1 Spastin expression dramatically increased the extent of LD-peroxisome contacts within cells, while depletion of M1 Spastin by siRNA significantly reduced LD-peroxisome contacts. Indeed, CLEM and FIB-SEM analysis showed that peroxisomes become tightly arrayed along the surface of the LDs upon M1 Spastin overexpression. We also found that M1 Spastin is highly enriched on LDs, but not on peroxisomes or significantly on ER, using its hairpin motif to insert into the lipid monolayer of LDs. By systematically exploring regions of M1 Spastin that promote LD-peroxisome tethering, we identified a PXI region of M1 Spastin for interacting with peroxisomes. The PXI region was sufficient to mediate LD-peroxisome association when appended to Spastin’s hairpin motif. Significantly, the PXI region in M1 Spastin forms a complex with ABCD1 in peroxisomes for LD-peroxisome tethering.

To address whether LD-peroxisome association mediated by M1 Spastin and ABCD1 enables directed FA transport between these organelles, we employed a FA pulse-chase labeling assay to measure FA transport. Cells with M1 Spastin depleted by siRNA had decreased FA transfer from LDs into peroxisomes, whereas those having increased M1 Spastin levels showed boosted transfer, implicating M1 Spastin as an enabler of LD-to-peroxisome FA trafficking. In exploring the mechanism underlying M1 Spastin-mediated FA trafficking, we discovered that M1 Spastin’s MIT domain, which interacts with ESCRT-III components, was necessary for FA trafficking. We further found that M1 Spastin’s MIT domain recruits the positive-curvature generating ESCRT-proteins IST1 and CHMP1B to LDs, and without this recruitment, transfer of FAs from LDs into peroxisomes was inhibited. These results indicated that tethering of LDs to peroxisomes and recruitment of IST1 and CHMP1B to LDs are both mediated by M1 Spastin and that these interactions were necessary for LD-to-peroxisome FA trafficking.

How might these events be conceptualized to explain the functions elicited by M1 Spastin at LD-peroxisome contact sites? We propose the following model (Fig. 7E). Tethering of LDs to peroxisome would be initiated when M1 Spastin’s PXI domain is made available to interact with ABCD1 on peroxisome membranes. We speculate that ATP hydrolysis by M1 Spastin’s ATPase domain may trigger a conformational change that favors PXI region and ABCD1 complex formation, since mutations in the ATPase domain disrupts LD-peroxisome contact formation. At LD-peroxisome contacts, FAs would be transferred from LDs to peroxisomes with the assistance of IST1 and CHMP1B. These proteins would serve to remodel the LD monolayer, potentially by their known ability to stabilize positively-curved membranes and to tubulate cellular membranes (McCullough et al., 2015).

This model raises many questions for future work. One issue is whether there are specific events that trigger the association of M1 Spastin with ABCD1 on peroxisomes. Our finding that the ATPase activity of M1 Spastin can regulate this interaction suggests it might be metabolically triggered by fluctuations in cellular ATP levels. Another question is how IST1 and CHMP1B function to facilitate FA trafficking. One crucial step for FA trafficking is FA release from LDs, which requires lipases to gain access to the lipids within LDs and is likely facilitated by increasing LD membrane curvature (Leff and Granneman, 2010). Given that IST1 and CHMP1B are capable of modifying lipid bilayers (McCullough et al., 2015), we speculate that IST1 and CHMP1B recruitment onto LDs by M1 Spastin could increase local membrane curvature by tubulating the lipid monolayer to facilitate FA trafficking. In yeast, high-curvature peroxisomal protrusions enriched in FA β-oxidation enzymes were found to extend into the lipid core of LDs at LD-peroxisome contacts (Binns et al., 2006). Therefore, altering membrane morphology at LD-peroxisome contacts seems to be a common mechanism to facilitate FA trafficking and metabolism throughout evolution. Because ESCRT-III proteins require VPS4 ATPase for assembly/disassembly cycles, one can ask whether VPS4 or possibly the ATPase activity from M1 Spastin is important for the remodeling of IST1 and CHMP1B at LD-peroxisome contacts to support FA trafficking.

This model may be instructive for understanding the pathogenesis of HSP diseases. It is increasingly clear that defects in FA metabolism are a common cause of HSP, and many of HSP-associated mutations affect proteins that impact FA allocation in LDs. For example, HSP-associated proteins involved in LD biogenesis such as Atlastin-1, REEP1 and Seipin are responsible for FA incorporation in LDs (Falk et al., 2014; Klemm et al., 2013; Renvoisé et al., 2016). Furthermore, HSP-associated lipase DDHD2 and its related proteins, DDHD1 and PNPLA6, may contribute to the release of FAs from LDs (Inloes et al., 2014; Rainier et al., 2008; Tesson et al., 2012). As demonstrated in this work, M1 Spastin and ABCD1 appear to have an important role in direct channeling of FA from LDs into peroxisomes. Together, these combined data suggest that rather than impacting disparate pathways associated with HSP, the mutations are all affecting the same pathway involving FA incorporation into LDs, FA release from LDs and delivery to peroxisomes. When this pathway is dysfunctional, as occurs in certain types of HSP as well as diseases associated with peroxisomal defects, LDs are unable to deliver FAs into peroxisomes, causing major downstream consequences for lipid detoxification, FA/lipid homeostasis and energy production.

## Materials and Methods

### Reagents

BODIPY 493/503, BODIPY 665/676, BODIPY 581/591 C11, cumene hydroperoxide, anti-CHMP1B antibody (PA5-44773), Dulbecco’s phosphate-buffered saline (DPBS), were purchased form ThermoFisher. 12-(7-Nitrobenzofurazan-4-ylamino)dodecanoic acid (NBD-C12), anti-ABCD1 antibody (ab197013), anti-PMP70 antibody (ab109448), anti-Catalase antibody (ab110292), anti-RFP antibody (ab124754) were obtained from Abcam. Oleic acid-BSA complex, nocodazole, anti-Spastin antibody (ABN368), anti-ACBD5 antibody (HPA012145), diethylumbelliferyl phosphate (DEUP) were purchased from Sigma-Aldrich. Anti-GAPDH antibody (2118) was purchased from Cell Signaling. Anti-IST1 antibody (GTX101972) was obtained from GeneTex. Formaldehyde and glutaraldehyde were obtained from Electron Microscopy Science. AP20187 (B/B Homodimerizer) was purchased from Takara. The recombinant Anti-M1 Spastin rabbit monoclonal antibody, Clone RM346 (Cat# 31-1232-00: RevMAb Biosciences USA, South San Francisco) was generated using RevMAb Biosciences’ proprietary B cell cloning technology against M1 specific sequences.

### Cell Culture and Transfection

HeLa cells (CCL-2) and U-2 OS (HTB-96) osteosarcoma cells were purchased from American Type Culture Collection (ATCC). HeLa and U-2 OS cells were maintained in EMEM (ATCC) and McCoy’s 5A medium (ATCC), respectively, with 10% fetal bovine serum (FBS, Corning) and 1X penicillin/streptomycin solution (Corning). MRC-5 fibroblasts were kindly provided by Dr. Evan Reid and cultured in DMEM (ThermoFisher) supplemented with 10% FBS and 1X penicillin/streptomycin solution.

DNA plasmids (15-50 ng) and siRNAs (~20 nM) were transfected into HeLa cells with TransIT-LT1 or TransIT-X2 reagent for 16-20 hours and TransIT-TKO reagent for 64-68 hours, respectively (Muris). For mEmerald-SKL or mCherry-SKL and siRNA cotransfection, HeLa cells were transfected simultaneously with 10 ng plasmid and 20 nM siRNA using both TransIT-LT1 and TransIT-TKO following manufacturer’s instruction. For plasmids and siRNA transfection in Figure 4, HeLa cells were first transfected with siRNA using TransIT-TKO for 48 hours and subsequently transfected with 15 ng mEmerald-SKL and 50 ng mApple-M1 Spastin using TransIT-X2 for ~16 hours. U-2 OS and MRC-5 cells were transfected with 15-50 ng DNA plasmids using TransIT-X2.

### Constructs and siRNAs

Sec61β-mEmerald, mEmerald-SKL and mCherry-SKL was provided by the Molecular Biology Core at Janelia Research Campus. Photoactivatable (PA)-GFP-SKL was generated by replacing the mEmerald portion of mEmerald-SKL with PA-GFP using AgeI and BsrGI restriction sites. Fluorescence protein-M1^1-92^ (FP-M1^1-92^) plasmids were generated by inserting a PCR fragment containing amino acid 1-92 of human M1 Spastin (NM_014946) into FP-C1 vectors using EcoRI and BamHI restriction sites. M1^1-92^-mApple plasmid was generated by inserting M1^1-92^ PCR fragment into mApple-N1 vector using BglII and HindIII restriction sites. 2xFKBP- mApple-M1^1-92^ was generated by inserting 2xFKBP PCR fragment into mApple-M1^1-92^ plasmid using NheI and AgeI sites. FP-M1 Spastin and FP-M87 Spastin plasmids were constructed the same way as FP-M1^1-92^ with PCR fragments containing amino acid 1-616 and 87-616 of M1 Spastin, respectively. We then introduced an M-to-A mutation at the 87^th^ amino acid in FP-M1 Spastin plasmids via site-direct mutagenesis to disrupt the second start codon and we used these constructs for subsequent cloning and mutagenesis. FP-M1 Spastin^Δ1-56^ was generated the same way as FP-M1^1-92^ with PCR fragments containing amino acid 57-616 of M1 Spastin. FP-M1 Spastin^HFDD^, FP-M1 Spastin^K388R^, and FP-M1 Spastin^E442Q^ were generated using site-directed mutagenesis. For mApple-M1^1-92^-197-328, PCR fragments containing M1 Spastin amino acids 197-328 was inserted into the C-terminus of FP-M1^1-92^ linearized by BamHI enzyme using In-Fusion cloning kit (Takara). DsRed-M1^1-92^-197-328 was constructed by replacing mApple in mApple-M1^1-92^-197-328 with DsRed2 using NheI and EcoRI restriction sites. mEmerald-GPAT4^152-208^ was constructed by inserting a PCR fragment containing amino acid 152-208 of human GPAT4 (NM_178819) into mEmerald-C1 vector using EcoRI and BamHI restriction sites. mEmerald-IST1 and mEmerald-CHMP1B were constructed by inserting PCR fragments of full-length IST1 (NM_001270975) and CHMP1B (NM_020412.4), respectively, into mEmerad-C1 vector using SalI and KpnI restriction sites. All oligonucleotides used in plasmid generation are listed in Table S1. Home-made siRNAs used in this study were generated by giardia dicer as previously described (Guiley et al., 2012; Liou et al., 2005). All oligonucleotides used in siRNA generation are listed in Table S2.

### Fluorescence Microscopy Imaging

All cells were grown and transfected on Lab-Tek II chambered #1.5 coverglasses (ThermoFisher) or MatTek dishes with #1.5 coverslip (MatTek Corporation; Ashland, MA). To label LDs, BODIPY 493/503 or BODIPY 665/676 was added to cells at 500 ng/ml 5 min or 16 hours prior to imaging, respectively, and was present during imaging. Confocal microscopy was performed on a custom-built Nikon Eclipse Ti microscope with a YOKOGAWA CSU-X1 Spinning Disk Confocal unit using SR Apo TIRF 100x/1.49 or Plan Apo λ 60x/1.40 objectives. Airyscan and structure illumination microscopy (SIM) imaging were performed using Plan Apo 63x/1.4 objective on ZEISS LSM 880 with Airyscan and ZEISS ELYRA Super-resolution Microscopy, respectively. Live-cell confocal and Airyscan experiments were conducted with cells incubated in phenol red-free medium at 37°C with 5% CO_2_ and humidified air. Photoactivation was performed on the custom-built Nikon microscope equipped with Bruker photoactivation module. HeLa cells overexpressing PA-GFP-SKL and mApple-M1 Spastin were subjected to 405 nm laser pulse at maximal intensity in 3.14-μm^2^ circular areas containing LDs labeled with BODIPY-665/676. For SIM experiments, transfected cells were washed with DPBS once and fixed with 4% formaldehyde and 0.1% glutaraldehyde for 20 min at room temperature. Fixed cells were then quenched with 100 mM glycine in DPBS followed by DPBS wash twice and subjected to SIM imaging at room temperature.

### Immunostaining

All procedures were performed at room temperature unless otherwise indicated, and all washing steps were done by DPBS for 5 min. HeLa cells transfected with mApple-M1 Spastin were rinsed with DPBS and fixed 4% formaldehyde and 0.1% glutaraldehyde for 20 min. Fixed cells were quenched with 100 mM glycine in DPBS, washed twice, and incubated with 0.3% Triton X-100 in DPBS for 20 min for permeabilization. Permeabilized cells were then blocked with 5% normal donkey serum (Jackson ImmunoResearch Laboratories, Inc.) in DPBS for 1 h followed by incubation with anti-catalase antibody (1:200 dilution) or anti-PMP70 antibody (1:200 dilution) in DPBS with 1% BSA at 4°C overnight. After three washes, the cells were incubated with fluorescent secondary antibody (1:2,000 in dilution) for 1 h. The stained samples were washed for three times and imaged with confocal microscopy at room temperature.

### Image Analysis

All image analyses were performed using ImageJ (National Institutes of Health) unless otherwise indicated. All intensity analyses were subjected to background subtraction. To obtain relative intensity profiles, the intensity values from different conditions were normalized to that at the first time point or in control groups. Co-localization of LDs and peroxisomes was quantified by measuring the Mander’s overlap coefficient of confocal maximal intensity projection (MIP) images using the JACoP plugin. For intensity analysis of 2xFKBP-mApple-M1^1-92^, regions of interest (ROIs) in LDs and the ER were generated using oval selection and polygon selection, respectively. Mean grey values of 2xFKBP-mApple-M1^1-92^ intensity in LDs were obtained and then averaged. For NBD-C12 intensity analyses in pulse-chase experiments, ROIs of LDs and peroxisomes were generated using their respective confocal MIP images. These ROIs were then applied to NBD-C12 confocal MIP images to obtain the mean grey values of NBD-C12 in LDs and peroxisomes. Similar analyses were applied to obtain NBD-C12 intensity in peroxisomes in transient pulse-chase assay with control cells or with M1 Spastin^K388R^ overexpressing cells. Due to the close association of LDs and peroxisomes in M1 Spastin or M1 Spastin^HFDD^ overexpressing cells, ROIs generated from peroxisome mask showed a substantial overlap with LDs. Thus, we manually selected peroxisome ROIs without overlapping LDs to analyze NBD-C12 intensity in transient pulse-chase assay with cells overexpressing M1 Spastin or M1 Spastin^HFDD^. For analyses of lipid peroxidation in LDs, ROIs of LDs were generated using oval selection. Mean grey values of ROIs from both green (Em 510 nm) and red (Em 590 nm) channels were extracted. Relative lipid peroxidation was derived from the ratio of Em 510 nm to Em 590 nm.

### Correlative Light-Electron Microscopy (CLEM)

HeLa cells grown on 25-mm photoetched coverslips (Electron Microscopy Sciences, Hatfield, PA) were transfected with 250 ng mApple-M1 Spastin and 100 ng mEmerald-SKL plasmids for ~16 hours in the presence of 15 μM oleic acid. Cells were then washed with DPBS and fixed with 4% FA, 0.1% GA 2mM CaCl_2_ in 0.08M sodium cacodylate buffer. Fixative was present during light microscopy image acquisition. M1 Spastin and peroxisome fluorescence images of cells of interest and their corresponding etched number on the coverslips were acquired by ZEISS LSM 880 with Airyscan. Subsequently, the fixative for light microscopy was removed and fresh 2% GA, 2 mM CaCl_2_ in 0.08 M sodium cacodylate buffer, pH 7.2 was added. Cells were kept in fixative at 4°C for 16 hours and postfixed in 2% osmium tetroxide-1.25% potassium ferrocyanide in 0.1M cacodylate buffer for 30 min followed by 2% osmium in the same buffer for another 30 min and processed for Epon embedding. Cells imaged by Airyscan were localized on the grid (imprinted in the Epon block). Ultrathin sections (60 nm) from the imaged cells were cut and post-stained with uranyl acetate/lead citrate and imaged in a Tecnai 12 electron microscope (FEI, Hillsboro, OR) operating at 80kV equipped with an Ultrascan 4000 digital camera (Gatan Inc, CA).

### Focused Ion Beam Scanning EM (FIB-SEM)

#### Sample preparation

HeLa cells grown on sapphire coverslips (3 mm diameter, 0.05 mm thickness, Nanjing Co-Energy Optical Crystal Co., Ltd) were transfected with mApple-M1 Spastin plasmid for ~16 hours in the presence of 15 μM oleic acid. Widefield fluorescence and DIC images were collected prior to high-pressure freezing for subsequent identification of cells of interest. The sapphire coverslips were clamped between two aluminium planchettes (Technotrade International, Manchester, NH) using 25% dextran as a filler. High-pressure freezing was performed in a Wohlwend Compact 01 high pressure freezer (Wohlwend GmbH, Switzerland), followed by freeze-substitution (FS). Coverslips were released from the planchettes and transferred to cryotubes containing FS medium (2% OsO_4_, 40mM imidazole, 40mM 1,2,4-triazole, 0.1% Uranyl acetate, and 4% water in acetone) under liquid N_2_. FS was performed using an automated FS machine (AFS2, Leica Microsystems, Buffalo Grove, IL) with FS schedule adapted from (Buser and Walther, 2008) as follows: −140°C to −90°C for 2 hours, −90°C for 24 hours, −90°C to 0°C for 12 hours, 0 to 22°C for 1 hour and 22°C for 90 min. Resin embedding was performed immediately after FS. Samples were removed from the FS machine, washed 3 times in anhydrous acetone for a total of 10 min and embedded in Eponate 12 as follows: Acetone/Eponate 12 (2:1) for 1 hour, Acetone/Eponate 12 (1:1) for 1 hour, Acetone/Eponate 12 (1:2) for 1 hour, Eponate 12 for 2 hours, Eponate 12 2 hours till overnight and Eponate 12 for 2 hours. Coverslips with cells side up were then placed in the slots of a flat embedding silicone mold filled with Eponate 12, and incubated at 60°C for 2 days to allow Eponate 12 polymerization. The epoxy was removed using a razor blade to expose the sapphire coverslip surface not containing cells. Coverslip was separated by sequential dipping the sample into liquid N_2_ and hot water. The exposed surface was immediately re-embedded in Durcupan, which helps reducing the streaks during FIB-SEM imaging (Xu et al., 2017). X-Ray of the entire re-embedded block was taken using XRadia 510 Versa micro X-Ray system (Carl Zeiss X-ray Microscopy, Inc., Pleasanton, CA). X-Ray images were then overlaid with fluorescence and DIC images to identify cells of interest. Sample block was then re-mounted using Durcupan on a copper stud (Xu et al., 2017) and trimmed using ultramicrotome (EM UC7, Leica Microsystems, Buffalo Grove, IL). Finally, the sample was sputter coated with 10nm of Au and 100nm of Carbon in a sputter-coating system (PECS 682, Gatan, Pleasanton, CA).

#### FIB-SEM imaging

FIB-SEM imaging was performed using a customized Zeiss Merlin crossbeam system previously described (Xu et al., 2017). The Zeiss Capella FIB column was repositioned at 90 degrees to the SEM column. The block face was imaged by a 2 nA electron beam with 1.2 keV landing energy at 500 kHz. The x-y pixel resolution was set at 8 nm. A subsequently applied focused Ga^+^ beam of 15 nA at 30 keV strafed across the top surface and ablated away 4 nm of the surface. The newly exposed surface was then imaged again. The ablation – imaging cycle continued about once every 30 seconds for one week. The sequence of acquired images formed a raw imaged volume, followed by post processing of image registration and alignment using a Scale Invariant Feature Transform (SIFT) based algorithm. The aligned stack was binned by a factor of 2 along z to form a final isotropic volume of 100 x 7 x 80 μm^3^ with 8 x 8 x 8 nm^3^ voxels, which can be viewed in arbitrary orientations. Organelle segmentation and 3D rendering were manually annotated using Amira software (ThermoFisher).

### Immunoprecipitation (IP) and Western Blotting

HeLa cells were cultured on 6-cm dishes and transfected with 1 μg of plasmid DNA for ~18 hours in the presence of 15 μM oleic acid. Cells were then washed with DPBS before lysis with IP lysis buffer (ThermoFisher) supplemented with 1x Halt protease inhibitor cocktail (ThermoFisher) on ice for 30 min. The lysates were subjected to centrifugation at 16,000 g for 15 min at 4°C, and the clear lysates (supernatants) were collected. The clear lysates were mixed with RFP-nAb agarose resins (Allele Biotechnology) and incubated with tumbling at 4°C for 4 h. The immunoprecipitated proteins were eluted with NuPAGE LDS sample buffer (ThermoFisher) after washing the RFP-nAb agarose resins twice with IP lysis buffer and twice with 10 mM Tris buffer containing 500 mM NaCl. To determine endogenous protein expression levels following siRNA treatment, cells grown in 12-well plate were transfected with siRNAs for ~70 hours and the cell lysates were collected as described above. The eluted proteins of IP experiments and lysates for siRNA experiments were analyzed by Western blotting. Quantification of IP was done by densitometry analysis using ImageJ.

### Statistical Analyses

Data were statistically analyzed by two-tailed *t* test or one-way ANOVA with Tukey’s multiple comparisons using GraphPad Prism (GraphPad Software; La Jolla, CA).

## Declaration of Interests

The authors declare no financial interests.

## Acknowledgements

We would like to thank Dr. Kevin McGowan and Jordan Towne in the Molecular Biology Core at Janelia Research Campus for plasmid preparation, Dr. Luke Lavis at Janelia Research Campus for providing JF Halo ligands, Dr. Evan Reid at Cambridge Institute for Medical Research, United Kingdom, for kindly providing MRC-5 cells, Rick Webb at University of Queensland, Australia, for advice on freeze-substitution, the Lippincott-Schwartz laboratory members for valuable discussions and technical assistance, and Victoria Custard for administrative assistance. This work was supported by HHMI Janelia Research Campus and the Intramural Research Program of the NINDS, NIH.

## Author Contributions

C.-L.C., C.B., and J.L.-S. conceived and designed the study. C.-L.C., M.S.I., H.A.P., C.S.X., D.R.P., G.S., and M.F. performed the experiments. C.-L.C., A.V.W., and M.S.I., analyzed the data. H.F.H. supervised the FIB-SEM experiment. C.-L.C. and J.L.-S. wrote the manuscript with the input from all co-authors. J.L.-S. supervised the project.

