## Supplemental Figures and Legends for "Spastin tethers lipid droplets to peroxisomes and directs fatty acid trafficking through ESCRT-III"

**Supplemental Figure Legends**

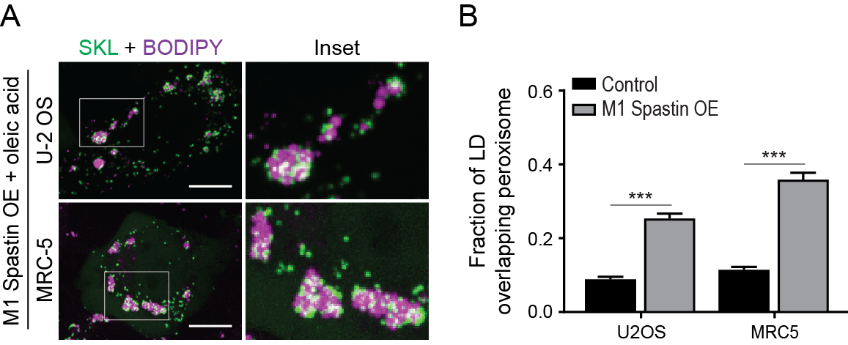

**Figure S1. M1 Spastin promotes LD-peroxisome contact formation in U2-OS and MRC-5 cells.**

(A) Association between BODIPY-665/676-labeled LDs and mEmerald-SKL-labeled peroxisomes in U-2 OS (top) and MRC-5 cells (bottom) expressing mApple-M1 Spastin in the presence of 300 μM oleic acid treatment. Representative confocal MIP images are shown. Scale bars, 10 µm.

(B) Quantification of fraction of LD overlapping with peroxisomes as described in (A). Means ± SEM are shown (23-34 cells from 3 independent experiments). ***, p < 0.001.

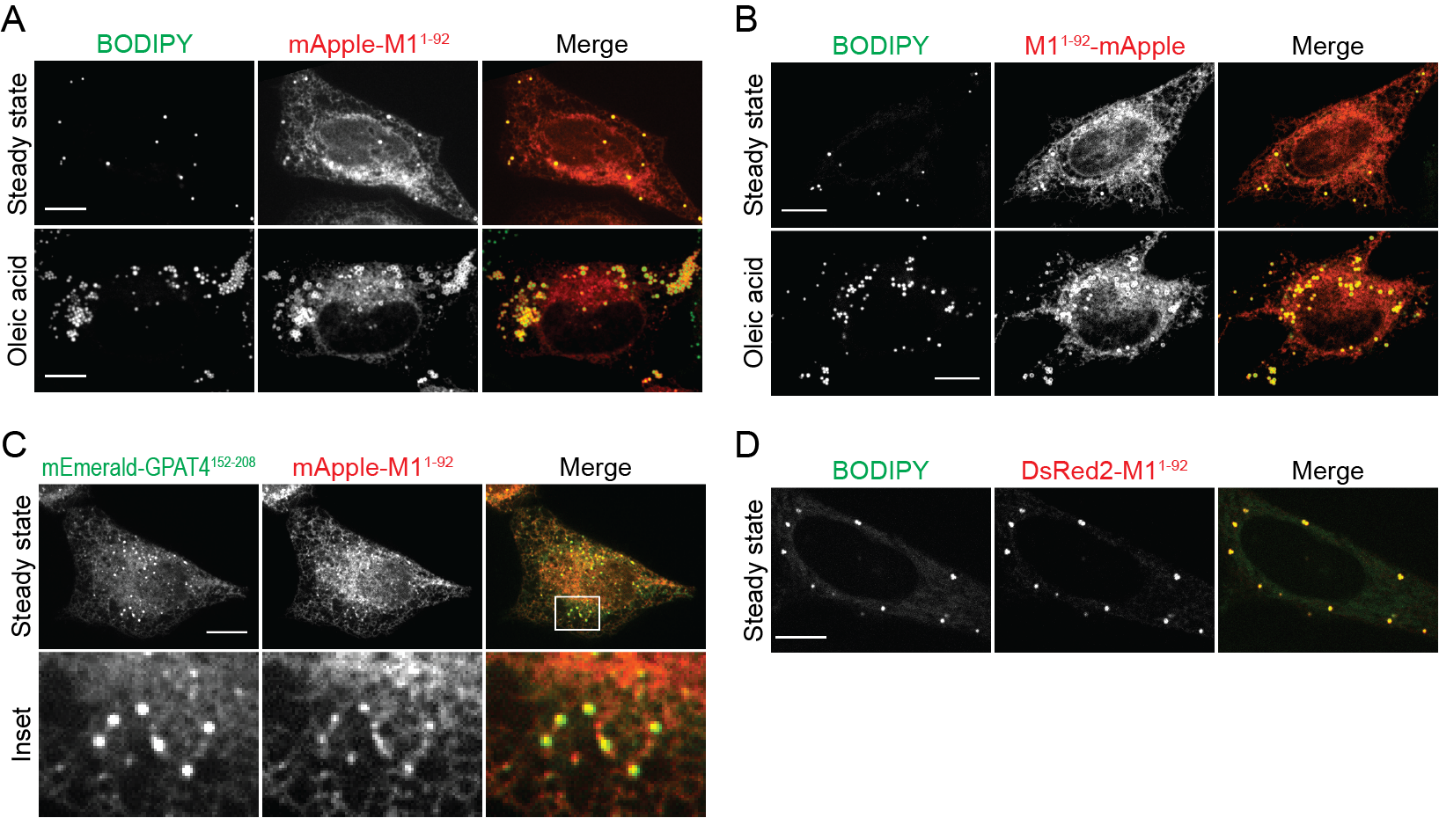

**Figure S2. M1 Spastin’s hairpin motif co-localizes with LDs.**

(A and B) mApple-M1^1-92^ (A) and M1^1-92^-mApple (B) display ER localization and co-localize with BODIPY-493/503-labeled LDs in HeLa cells in steady state (top) or following 300 μM oleic acid treatment for 16 hours (bottom). Representative confocal images are shown. Scale bar, 10 µm.

(C) mApple-M1^1-92^ co-localizes with mEmerald-GPAT4^152-208^ in steady-state HeLa cells monitored by confocal microscopy. Scale bar, 10 µm.

(D) DsRed2-M1^1-92^ co-localizes with BODIPY-665/676-labeled LDs in HeLa cells in steady state. Representative confocal images are shown. Scale bar, 10 µm.

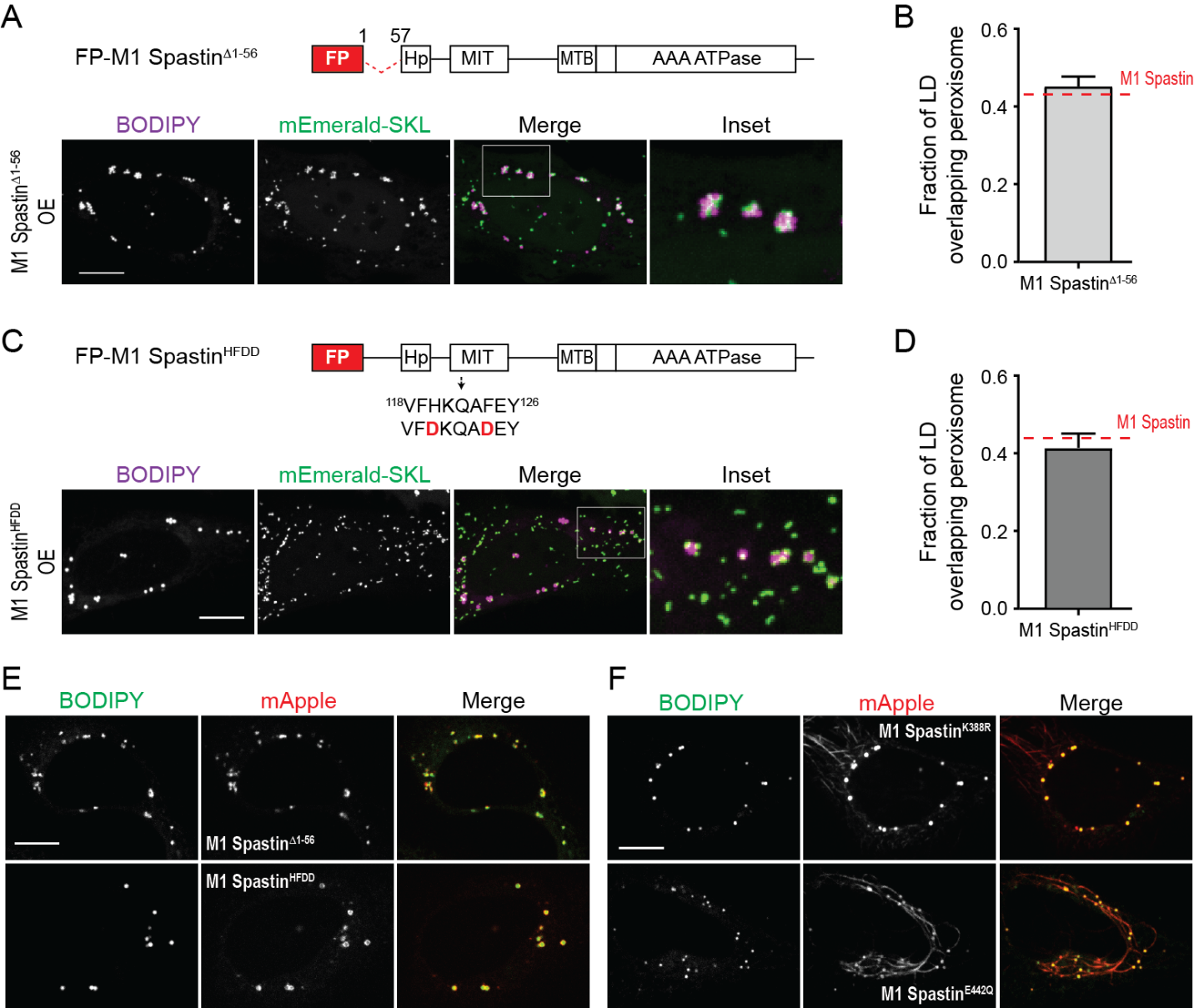

**Figure S3. Subcellular localization M1 Spastin mutants and synthetic constructs in respective to LDs and peroxisomes.**

(A) Diagrams of FP-M1 Spastin^Δ1-56^ and the association between BODIPY-665/676-labeled LDs and mEmerald-SKL-labeled peroxisomes in Hela cells expressing mApple- M1 Spastin^Δ1-56^. Representative confocal MIP images are shown. Scale bar, 10 µm.

(B) Quantification of fraction of LD overlapping with peroxisomes as described in (A) and in cells expressing mApple-M1 Spastin (red dashed line). Mean ± SEM is shown (28 cells from 3 independent experiments).

(C) Diagrams of FP-M1 Spastin^HFDD^ and the association between BODIPY-665/676-labeled LDs and mEmerald-SKL-labeled peroxisomes in Hela cells expressing mApple- M1 Spastin^HFDD^. Representative confocal MIP images are shown. Scale bar, 10 µm.

(D) Quantification of fraction of LD overlapping with peroxisomes as described in (C) and in cells overexpressing mApple-M1 Spastin (red dashed line). Mean ± SEM is shown (33-34 cells from 3 independent experiments).

(E) Co-localization of BODIPY-665/676-labeled LDs and mApple-M1 Spastin^Δ1-56^ or mApple-M1 Spastin^HFDD^ in HeLa cells. Representative confocal images are shown. Scale bar, 10 µm.

(F) Co-localization of BODIPY-665/676-labeled LDs and mApple-M1 Spastin^K388R^ or mApple-M1 Spastin^E442Q^ in HeLa cells. Representative confocal images are shown. Scale bar, 10 µm.

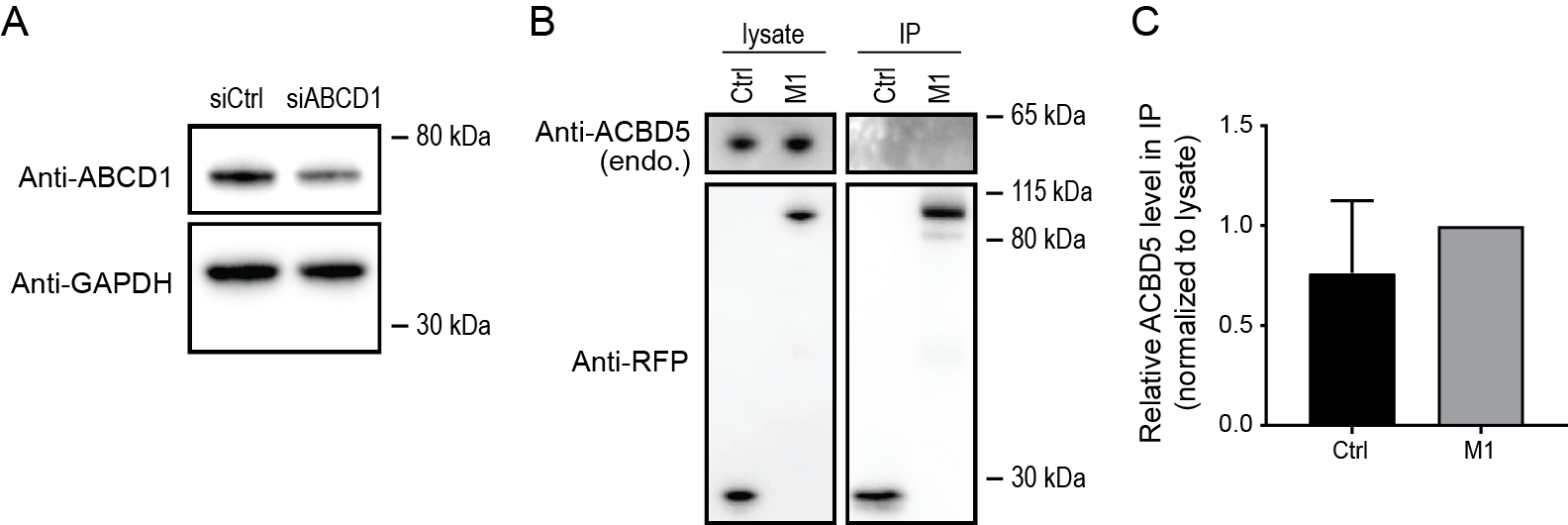

**Figure S4. Endogenous ABCD1 protein levels and minimal interaction between M1 Spastin and ACBD5.**

(A) Endogenous ABCD1 protein levels detected by Western blotting using anti-ABCD1 antibody (top) in HeLa cells transfected with siCtrl or siABCD1. GAPDH protein levels serve as a loading control (bottom).

(B) IP of ACBD5 in Hela cells transfected with mApple-C1 vector (Ctrl) and mApple-M1 Spastin (M1). Protein levels of endogenous ACBD5 and overexpressed constructs in cell lysates and IP were assessed by Western blotting using antibodies against ACBD5 and RFP.

(C) Quantification of relative ACBD5 level in the IP as described in (B). The value of M1 was set as 1. Means ± SEM from 5 independent IP experiments are shown.

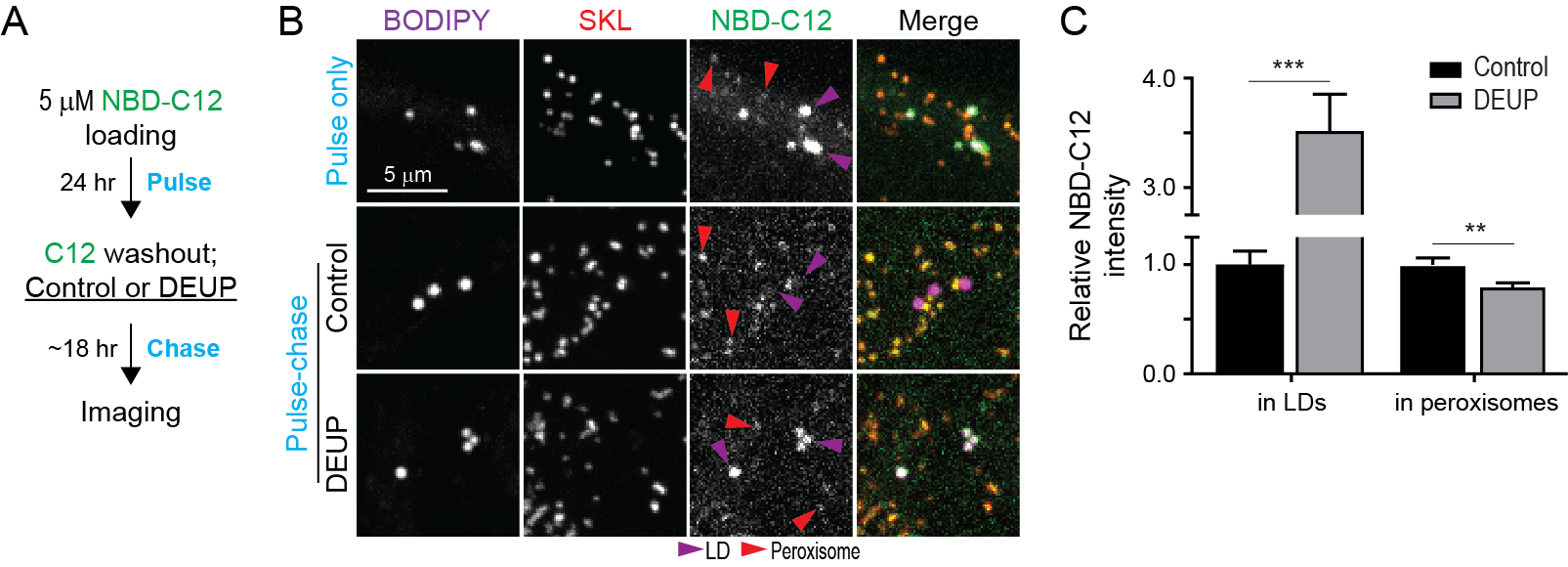

**Figure S5. Monitoring LD-to-peroxisome FA trafficking using NBD-C12.**

(A) Schematic diagram representing fluorescence FA pulse-chase assay using NBD-C12 with DEUP treatment.

(B) Distribution of NBD-C12 in BODIPY-665/676-labeled LDs and mCherry-SKL-labeled peroxisomes after pulse (top) and after pulse-chase in control cells (middle) or in cells treated with 150 μM DEUP (bottom). Magenta and red arrow heads indicate LDs and peroxisomes, respectively. Representative confocal MIP images are shown.

(C) Relative NBC-C12 intensity in LDs or peroxisomes as described in (B). Means ± SEM are shown (26-27 cells from 3 independent experiments). **, p < 0.01; ***, p < 0.001.

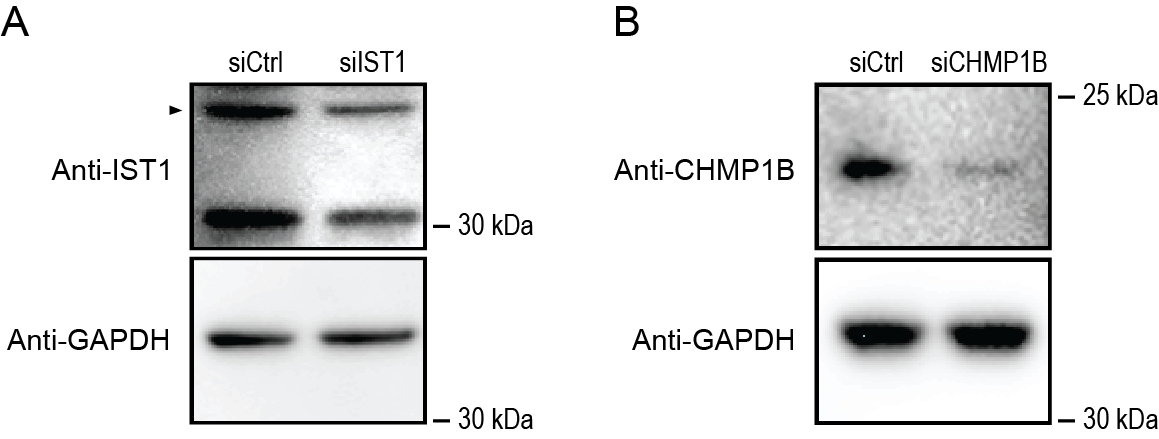

**Figure S6. Detecting endogenous IST1 and CHMP1B protein levels.**

(A) Endogenous IST1 protein levels detected by Western blotting using anti-IST1 antibody (top) in HeLa cells transfected with siCtrl or siIST1. GAPDH protein levels serve as a loading control (bottom).

(B) Endogenous CHMP1B protein levels detected by Western blotting using anti-CHMP1B antibody (top) in HeLa cells transfected with siCtrl or siCHMP1B. GAPDH protein levels serve as a loading control (bottom).

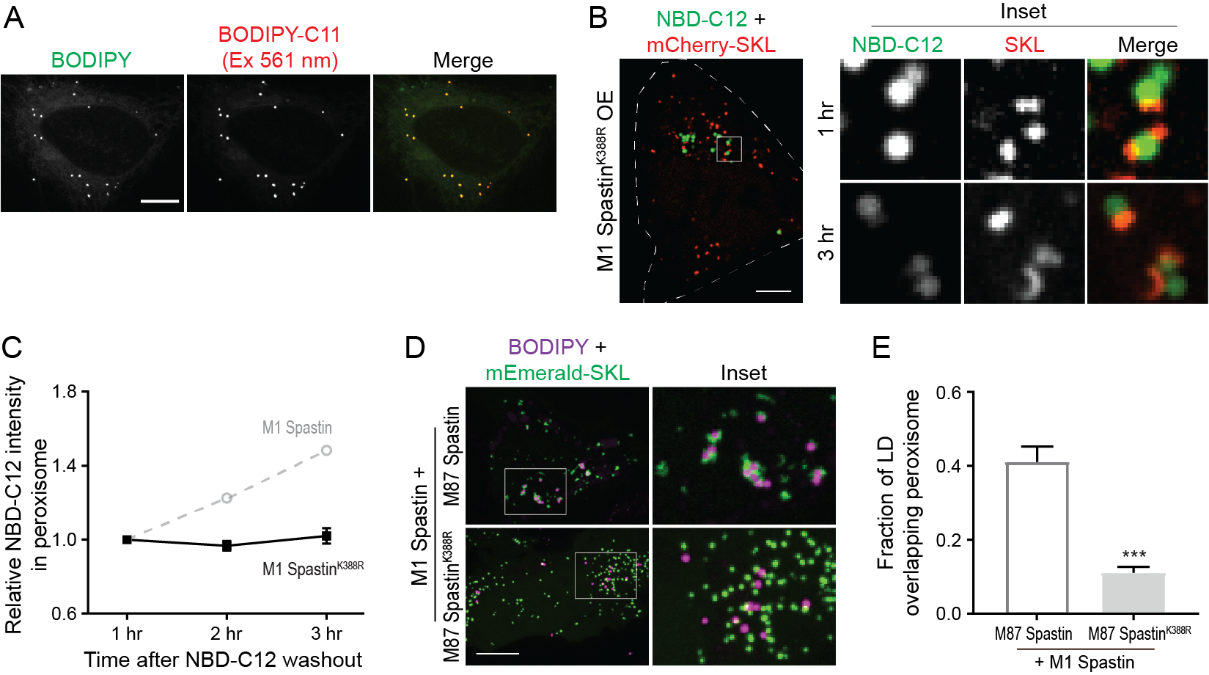

**Figure S7. Pathogenic K388R mutation redistributes M1 Spastin and disrupts LD-peroxisome contact formation.**

(A) Co-localization of BODIPY-C11 (imaged at Ex 561 nm) and BODIPY-labeled LDs in Hela cells. Representative confocal MIP images are shown. Scale bar, 10 µm.

(B) Distribution of NBD-C12 in Halo-M1 Spastin^K388R^ and mCherry-SKL overexpressing HeLa cells monitored by confocal microscopy. Scale bar, 10 µm.

(C) Relative NBC-C12 intensity in peroxisomes as described in (B). Means ± SEM are shown (15 cells from 3 independent experiments). Gray dashed line is a replica from Figure 5F.

(D) Association between BODIPY-665/676-labeled LDs and mEmerald-SKL-labeled peroxisomes in Hela cells co-expressing mApple-M1 Spastin and mTagBFP2-M87 Spastin or mTagBFP2-M87 Spastin^K388R^. Representative confocal MIP images are shown. Scale bar, 10 µm.

(E) Quantification of the fraction of LD overlapping with peroxisomes as described in (C). Means ± SEM are shown (13-22 cells from 2 independent experiments). ***, p < 0.001.

**Supplemental Video 1.** **Volumetric, Spastin driven peroxisome-LD contacts.**

Animation of 3D rendering in Figure 1G. Movie begins by sectioning through the *z* axis, then repeating with the annotated LDs (red) and peroxisomes (cyan), and once more with the 3D rendering of the LD and peroxisome segmentations. The 3D renderings are rocked about the *x*-axis. A 360-degree rotation with all the encompassing organelles in the volume follows (Green: Mitochondria, Blue: ER, Gray: Plasma Membrane, Purple: Endosome, Orange: Multi-Vesicular Body). Bounding box is 1.2 µm x 1.2 µm x 1.6 µm.
