## Supplementary material for "Spastin tethers lipid droplets to peroxisomes and directs fatty acid trafficking through ESCRT-III": Table S1

| **Table S1. Oligonucleotides for plasmid generation.** | |
| --- | --- |
| **Name** | **Sequence (5’-3’)** |
| M1 Spastin F | atatgaattctccgggtggac (EcoRI) |
| Spastin R | atcgggatccttaaacagtggtatctccaaagtc (BamHI) |
| FP-M1^1-92^ F | Same as M1 Spastin F |
| FP-M1^1-92^ R | atcgggatccttagctcctcttggctgccatg (BamHI) |
| M1^1-92^-FP F | atcgagatctcaccatgaattctccgggtggac (BglII) |
| M1^1-92^-FP R | atcgaagcttgctcctcttggctgccatg (HindIII) |
| 2xFKBP F | atcggctagcaccatgggtgctctcgagggagtgca (NheI) |
| 2xFKBP R | atcgaccggtagtgcactgcctccagctga (AgeI) |
| M1 Spastin^K388R^ F | ccacctgggaatgggaggacaatgctggctaaagcag |
| M1 Spastin^K388R^ R | tttagccagcattgtcctcccattcccaggtggac |
| M1 Spastin^E442Q^ F | ataatttttatagatcaagttgatagccttttgtgt |
| M1 Spastin^E442Q^ R | aaaggctatcaacttgatctataaaaattatagaaggttgaa |
| M1 Spastin^Δ1-56^ F | atcggaattctctgtttgtaggcttcgcg (EcoRI) |
| M1 Spastin^Δ1-56^ R | Same as Spastin R |
| M1 Spastin^HFDD^ F | cgcgtccgagtcttcgacaaacaggccgacgagtacatctccattgccc |
| M1 Spastin^HFDD^ R | aatggagatgtactcgtcggcctgtttgtcgaagactcggacgcgctcg |
| M1^1-92^-197-328 F | gcagccaagaggagcaagatgcaaccagttttgcc |
| M1^1-92^-197-328 R | ttatctagatccggtggatccttactatataaggttagcaaggttgct |
| GPAT4^152-208^ F | atcgaattctaacttccagtacatcagcctt (EcoRI) |
| GPAT4^152-208^ R | atcgggatccttagaactccttaaacctcccatt (BamHI) |
| IST1 F | atcggtcgacctgggctctggatttaaag (SalI) |
| IST1 R | atcgggtaccctatgttttctttttcagctcttc (KpnI) |
| CHMP1B F | atcggtcgactctaacatggagaaacacctg (SalI) |
| CHMP1B R | atcgggtacctcacacttgatcccgaagg (KpnI) |
