## Supplementary material for "Spastin tethers lipid droplets to peroxisomes and directs fatty acid trafficking through ESCRT-III": Table S2

| **Table S2. Oligonucleotides for siRNA generation.** | |
| --- | --- |
| **Name** | **Sequence (5’-3’)** |
| siControl F | taatacgactcactatagggtttgccttcctgtttttgct |
| siControl R | taatacgactcactatagggttgccgggaagctagagtaa |
| siSpastin F | taatacgactcactatagggtttaaggccttgccttgatg |
| siSpastin R | taatacgactcactatagggtttaaggccttgccttgatg |
| siABCD1 F | taatacgactcactatagggtggagcccacaaagtctacc |
| siABCD1 R | taatacgactcactatagggggcttggtcaggttggagta |
| siIST1 F | taatacgactcactatagggccaaaatcctggtggagaga |
| siIST1 R | taatacgactcactatagggtccgggaaagatcatcaaag |
| siCHMP1B F | taatacgactcactatagggcggccaaagaactgagtag |
| siCHMP1B R | taatacgactcactatagggtcacacttgatcccgaagg |
